# A distinctive evolution of alveolar T cell responses is associated with clinical outcomes in unvaccinated patients with SARS-CoV-2 pneumonia

**DOI:** 10.1101/2023.12.13.571479

**Authors:** Nikolay S. Markov, Ziyou Ren, Karolina J. Senkow, Rogan A. Grant, Catherine A. Gao, Elizabeth S. Malsin, Lango Sichizya, Hermon Kihshen, Kathryn A. Helmin, Milica Jovisic, Jason M. Arnold, Xóchitl G. Pérez-Leonor, Hiam Abdala-Valencia, Suchitra Swaminathan, Julu Nwaezeapu, Mengjia Kang, Luke Rasmussen, Egon A. Ozer, Ramon Lorenzo-Redondo, Judd F. Hultquist, Lacy M. Simons, Estefany Rios-Guzman, Alexander V. Misharin, Richard G. Wunderink, G.R. Scott Budinger, Benjamin D. Singer, Luisa Morales-Nebreda, The NU SCRIPT Study Investigators

## Abstract

Pathogen clearance and resolution of inflammation in patients with pneumonia require an effective local T cell response. Nevertheless, local T cell activation may drive lung injury, particularly during prolonged episodes of respiratory failure characteristic of severe SARS-CoV-2 pneumonia. While T cell responses in the peripheral blood are well described, the evolution of T cell phenotypes and molecular signatures in the distal lung of patients with severe pneumonia caused by SARS-CoV-2 or other pathogens is understudied. Accordingly, we serially obtained 432 bronchoalveolar lavage fluid samples from 273 patients with severe pneumonia and respiratory failure, including 74 unvaccinated patients with COVID-19, and performed flow cytometry, transcriptional, and T cell receptor profiling on sorted CD8^+^ and CD4^+^ T cell subsets. In patients with COVID-19 but not pneumonia secondary to other pathogens, we found that early and persistent enrichment in CD8^+^ and CD4^+^ T cell subsets correlated with survival to hospital discharge. Activation of interferon signaling pathways early after intubation for COVID-19 was associated with favorable outcomes, while activation of NF-κB-driven programs late in disease was associated with poor outcomes. Patients with SARS-CoV-2 pneumonia whose alveolar T cells preferentially targeted the Spike and Nucleocapsid proteins tended to experience more favorable outcomes than patients whose T cells predominantly targeted the ORF1ab polyprotein complex. These results suggest that in patients with severe SARS-CoV-2 pneumonia, alveolar T cell interferon responses targeting structural SARS-CoV-2 proteins characterize patients who recover, yet these responses progress to NF-κB activation against non-structural proteins in patients who go on to experience poor clinical outcomes.

## Introduction

Severe pneumonia due to SARS-CoV-2 is responsible for the bulk of the acute morbidity and mortality caused by the COVID-19 pandemic (1). Patients with severe SARS-CoV-2 pneumonia experience prolonged durations of respiratory failure when compared to patients with pneumonia secondary to other pathogens, increasing the risk of secondary bacterial pneumonia and other complications of critical illness (2, 3). During the pandemic, we used a systems biology approach that included data generated from bronchoalveolar lavage (BAL) sampling of the lungs of patients with severe pneumonia to suggest that spatially localized inflammatory circuits between alveolar macrophages and T cells sustain prolonged inflammation in patients with SARS-CoV-2 pneumonia (4). This model, which has since been confirmed by others (5-8), highlights the unusual importance of alveolar T cells in the immune response to SARS-CoV-2. Nevertheless, longitudinal data describing T cell responses to SARS-CoV-2 has been limited to analysis of peripheral blood samples (9-13). These studies suggest that a coordinated response between interferon-producing innate and adaptive immune cells drives viral clearance in patients with less severe COVID-19. In contrast, older patients with more severe disease exhibit excessive inflammatory immune cell activation and inadequate type I interferon production. Whether these responses reflect those in the alveolar space is not known. Furthermore, whether the T cell responses in the alveolus are unique to COVID-19 or are common to pneumonia irrespective of pathogen is also unknown, as most studies have compared patients with COVID-19 to healthy controls. These questions take on added importance as the pandemic wanes and scientists and clinicians question whether therapies shown to benefit patients with SARS-CoV-2 pneumonia will be effective in patients with pneumonia secondary to other pathogens.

We examined alveolar T cell responses in 432 BAL fluid samples collected serially from 273 patients with severe pneumonia, 74 of them with SARS-CoV-2 pneumonia, all of whom developed respiratory failure requiring mechanical ventilation. All of these samples were collected before effective vaccines for SARS-CoV-2 were produced. Compared to similarly ill patients with non-COVID-19 etiologies of pneumonia and respiratory failure, we found that abundance of T cell subsets expressing interferon-stimulated genes and targeting structural SARS-CoV-2 proteins (Spike and Nucleocapsid) were associated with survival, whereas a T cell activation profile dominated by a TNF-α/NF-κB inflammatory signature and enrichment in SARS-CoV-2 ORF1ab antigen specificity was associated with hospital mortality in patients with severe SARS-CoV-2 pneumonia. These findings suggest a unique pattern and evolution of T cell responses associated with severe SARS-CoV-2 infection in the alveolar space that may drive clinical outcomes (**Graphical Abstract**).

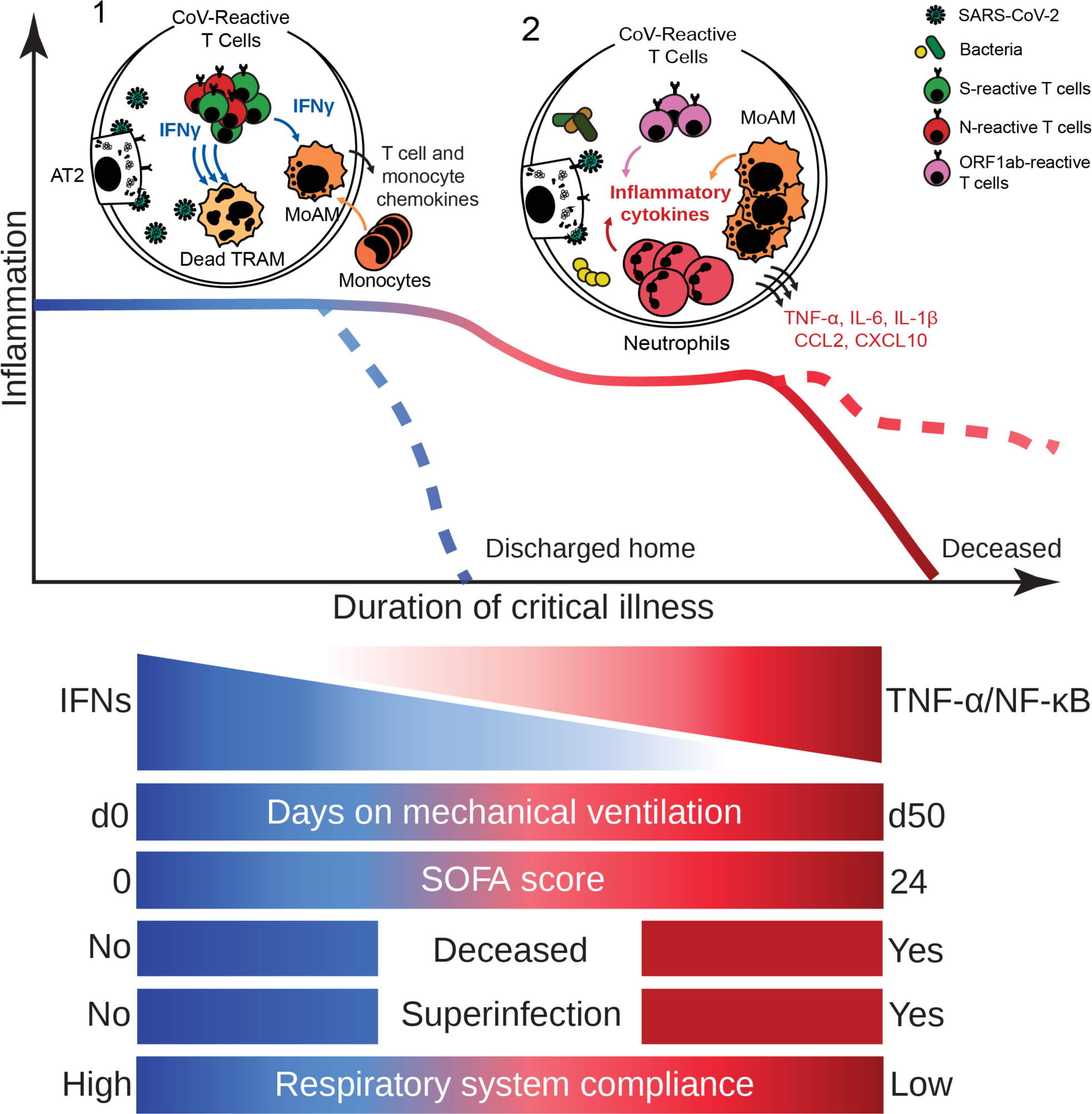
Graphical Abstract.

## Results

### Cohort demographics

The Successful Clinical Response in Pneumonia Therapy (SCRIPT) study is a prospective, single-center observational cohort study of mechanically ventilated patients who undergo lower respiratory tract sampling with at least one BAL procedure to evaluate suspected pneumonia as part of routine clinical care. To evaluate the alveolar T cell response to severe pneumonia, we focused on clinical BAL fluid samples obtained from June 2018 through August 2020 from patients with known or suspected pneumonia and respiratory failure requiring mechanical ventilation who consented to enroll in SCRIPT. During this time period, 273/337 (81%) participants enrolled in SCRIPT had at least one BAL fluid sample that underwent flow cytometry analysis; we identified these 273 patients as the cohort for this study (**Supplemental Figure 1A**). Of these 273 participants, 33 were adjudicated to have a non-pneumonia etiology for their respiratory failure (non-pneumonia controls, most often due to aspiration, cardiogenic pulmonary edema, or atelectasis). The remainder were diagnosed with pneumonia: 133 with pneumonia due to a bacterial pathogen (other pneumonia), 74 with SARS-CoV-2 pneumonia with or without bacterial superinfection (COVID-19), and 33 with pneumonia due to another virus with or without bacterial superinfection (other viral pneumonia). Some of these patients have been described in other publications (4, 14, 15). None of the patients had received SARS-CoV-2 vaccines, as the study period preceded their availability. Cohort demographics and clinical characteristics are shown in **Supplemental Table 1**, and some de-identified clinical information from this cohort has been published on PhysioNet (16). Demographics, body mass index, and the severity of illness as measured by Sequential Organ Failure Assessment (SOFA) score and Acute Physiology Score (APS) were similar across groups. Patients with COVID-19 had nominally lower rates of common comorbidities than the overall cohort. The percentages of samples obtained from study participants with viral pneumonia that also contained a bacterial superinfection were 46.1% for SARS-CoV-2 pneumonia and 38.9% for other viral pneumonia (*p* = 0.603, Fisher exact test). As most of these samples were collected before publication of the first randomized controlled trial that demonstrated efficacy of corticosteroids in patients with COVID-19 (the RECOVERY trial) (17), the administration of corticosteroids was not systematically guided by diagnosis. One-third of the cohort was received in external transfer from an outside hospital.

### Cellular composition and T cell immunophenotype of BAL fluid

We obtained 546 BAL fluid samples from the 337 patients enrolled in SCRIPT from June 2018 through August 2020. Partial analysis of flow cytometry data from a subset of the samples included in this study were previously published (4). Flow cytometry data were available in 432 (79.1%) of the samples (**Figure 1A** and **Supplemental Figure 1A-B).** (4) Hierarchical clustering performed on the normalized abundance of BAL fluid cell populations revealed a distinctive enrichment in T cells and monocytes in patients with COVID-19, irrespective of bacterial superinfection status (**Figure 1B** and **Supplemental Figure 2A-F**). Macrophages were enriched in non-pneumonia control samples, while neutrophils were higher in the other pneumonia group when compared with other groups (**Supplemental Figure 2G-H**). Compared to samples obtained from patients with COVID-19 ≤48 hours following intubation (early), the proportions of monocytes, CD3^+^ T cells, and T cell subsets (CD4^+^ T cells, CD8^+^ T cells, and Treg cells) were lower and the proportion of neutrophils was higher in BAL fluid samples collected from patients with COVID-19 >48 hours following intubation (late) (**Figure 1C-D** and **Supplemental Figure 2I-M**, **N-O, P, and S**). The proportion of macrophages was higher only in early samples from the non-pneumonia control group when compared with early samples from the other pneumonia group (**Supplemental Figure 2Q-R**).

**Figure 1.**
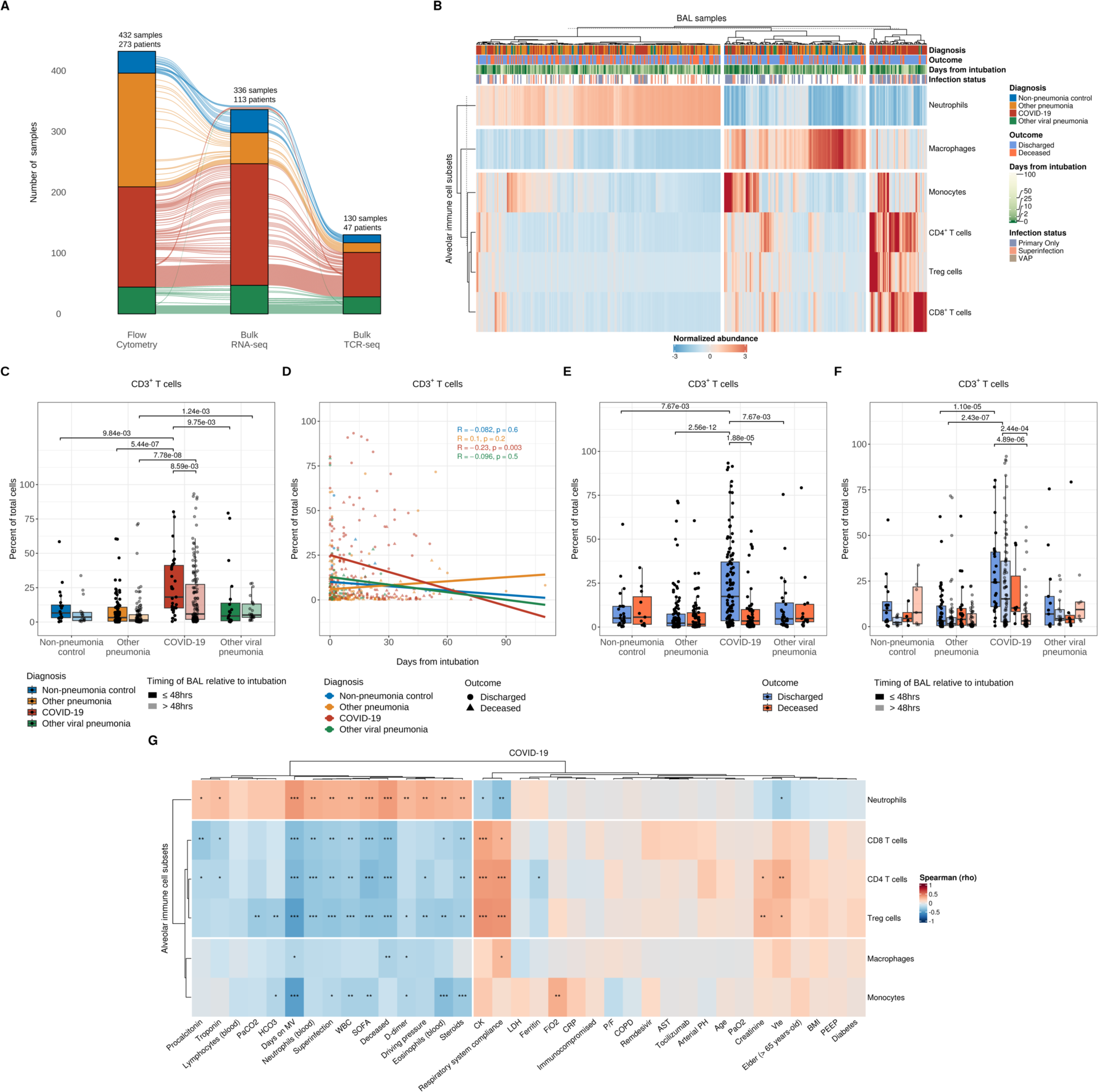
Alveolar T cell enrichment is associated with clinical outcomes in patients with severe SARS-CoV-2 pneumonia. **(A)** Alluvial diagram depicting multi-step analysis of BAL fluid samples with flow cytometry, bulk RNA-sequencing, and bulk TCR-sequencing in patients from non-pneumonia control, other pneumonia, COVID-19, and other viral pneumonia groups. The number of analyzed samples from unique SCRIPT-enrolled patients are illustrated at the top of each stratum. **(B)** Hierarchical clustering of flow cytometry analysis of alveolar immune cell subset composition from all samples. Each column represents a BAL sample and headers are color-coded by diagnosis, binary outcome (whether a given patient was discharged or died during hospitalization), duration of mechanical ventilation (blanks indicate chronically ventilated patients), and infection status (presence or absence of bacterial superinfection in patients with COVID-19 or other viral pneumonia). The VAP (ventilator-associated pneumonia) flag designates samples from non-pneumonia controls or patients with COVID-19 or other viral pneumonia who cleared the virus and then developed a bacterial pneumonia. Samples were clustered using Euclidean distance and Ward’s minimum variance linkage method. **(C)** Comparison of CD3^+^ T cell percentage between early (≤48 hours following intubation) and late (>48 hours following intubation) samples (*q* < 0.05, pairwise Wilcoxon rank-sum tests with FDR correction). **(D)** Correlation analysis between the percentage of alveolar CD3^+^ T cells and duration of mechanical ventilation with Pearson correlation coefficient. **(E-F)** Comparison of percent of alveolar CD3^+^ T cells between patients who were discharged from the hospital or died during their hospital course (E) and between early and late samples (F) (*q* < 0.05, pairwise Wilcoxon rank-sum tests with FDR correction). **(G)** Correlation analysis between the percentage of alveolar immune cell subsets and clinical, physiologic, and laboratory variables with Spearman rank correlation coefficient and FDR correction (*q* < 0.05 [*], *q* < 0.01 [**] and *q* < 0.001 [***]). Abbreviations: PaCO2 (partial arterial carbon dioxide pressure), HCO3 (bicarbonate), Days on MV (days on mechanical ventilation), SOFA (Sequential Organ Failure Assessment), WBC (peripheral white blood cells), CK (creatinine kinase), Vte (minute ventilation), LDH (lactate dehydrogenase), FiO2 (fraction of inspired oxygen), CRP (C-reactive protein), PEEP (positive end-expiratory pressure), BMI (body mass index), AST (aspartate aminotransferase), PaO2 (partial arterial oxygen pressure), P/F (ratio of partial arterial oxygen pressure to fraction of inspired oxygen), COPD (chronic obstructive pulmonary disease).

We next examined the association between hospital mortality and the proportion of immune cell subsets present in BAL fluid. Consistent with previous studies of BAL fluid or endotracheal tube washes from patients with severe SARS-CoV-2 pneumonia (18, 19), the abundance of CD3^+^ T cells—including CD4^+^ T cells, CD8^+^ T cells, and Treg cells—was positively associated with survival to hospital discharge (**Figure 1E** and **Supplemental Figure 3A, E and I**). BAL fluid T cell abundance was not associated with outcomes in patients with pneumonia secondary to other viruses or bacteria, suggesting a unique pathobiology in patients with SARS-CoV-2 pneumonia. In contrast to an analysis of endotracheal tube washes (19), we observed a positive association between survival and the overall abundance of macrophages, but not monocytes, in patients with COVID-19 (**Supplemental Figure 3C and G**), perhaps reflecting differences between endotracheal wash fluid and BAL fluid.

Changes in BAL fluid cell type composition over the course of disease were associated with clinical outcomes. Indeed, in patients with COVID-19, persistence of T cells in BAL fluid over time was associated with survival to hospital discharge, whereas a decrease in T cell and macrophage abundance was associated with mortality (**Figure 1F** and **Supplemental Figure 3B, D, F, H, and J**). In contrast, we found an inverse association between BAL fluid neutrophilia and survival to hospital discharge (**Supplemental Figure 3K**). Patients with COVID-19 who died in the hospital had greater neutrophilia in late samples than those who survived hospitalization (**Supplemental Figure 3L**), possibly reflecting the higher risk of late ventilator-associated pneumonia (VAP) in COVID-19 compared with pneumonia due to other causes (14, 15, 20) (see **Supplemental Figure 2P**). Consistent with this hypothesis, neutrophilic enrichment positively correlated with inflammatory markers (procalcitonin, troponin, D-dimer), steroid administration, severity of illness (SOFA score), driving pressure, duration of mechanical ventilation, presence of superinfection, and hospital mortality in patients with COVID-19 (**Figure 1G**), but not in patients with other causes of pneumonia and respiratory failure (**Supplemental Figure 4A-C**). In contrast, T cell enrichment was associated with favorable clinical parameters, including respiratory system compliance, in patients with COVID-19. The resolution status of the pneumonia episode based on expert clinical adjudication (21) was linked to mortality in patients with severe SARS-CoV-2 pneumonia in our cohort (**Supplemental Figure 3M-N**), similar to our findings in the larger cohort of patients with severe pneumonia enrolled in the SCRIPT study (15).

Expression of CD127 (the interleukin-7 receptor) and HLA-DR on peripheral blood T cells has been associated with outcomes in SARS-CoV-2 pneumonia, with CD127 correlating with less severe disease and better outcomes and HLA-DR correlating with worsened disease severity and poorer outcomes (9, 22, 23). Consistent with these observations, we found that expression of CD127 on CD8^+^ T cells in the alveolar space negatively correlated with severity of illness measured by SOFA score, while HLA-DR expression positively correlated with the inflammatory marker D-dimer among other variables (**Supplemental Figure 4D**). In summary, flow cytometry analysis of BAL fluid revealed enrichment in T cell subsets among patients with COVID-19 that predicted favorable clinical outcomes, including lower severity of illness, shorter duration of mechanical ventilation, and less hospital mortality.

### Transcriptional profiling of T cell subsets reveals a distinctive immune cell activation profile associated with clinical outcomes in patients with severe SARS-CoV-2 pneumonia

We flow cytometry sorted CD3ε^+^ T cell subsets into CD8^+^ T cells, CD4^+^ Treg cells (CD25^hi^CD127^lo^), and non-Treg CD4^+^ T cells (referred to here as CD4^+^ T cells) and analyzed samples with sufficient quantities of high-quality RNA using bulk RNA-sequencing (336 samples from 113 patients) (see **Figure 1A** and **Supplemental Figure 1B**). This subset largely resembled the larger cohort (**Supplemental Figure 5A-F**). We identified differentially expressed genes in CD8^+^ and CD4^+^ T cells (975 and 865 differentially expressed genes, respectively [FDR *q* < 0.05]) between patient groups. *K*-means clustering with *K* = 2 revealed genes in both CD8^+^ and CD4^+^ T cells that distinguished patients with COVID-19 from other groups (**Figure 2A**, **Figure 3A**, and **Supplemental Files 1-4**). In CD8^+^ T cells, Cluster 1 contained genes involved in cell proliferation, monocyte and T cell migration, tissue residency, and immune cell inhibition (**Supplemental Figure 6A**). Cluster 1 in CD4^+^ T cells contained genes associated with cell proliferation, immune cell activation, co-inhibitory molecules, markers of tissue resident-memory/effector-memory T cells, and monocyte and B cell chemoattractants (**Supplemental Figure 7A**). In both CD8^+^ and CD4^+^ T cells, genes in Cluster 2 (mostly downregulated in COVID-19) were associated with a resting or quiescent T cell program characteristic of naïve or central-memory T cells. We then examined gene ontology (GO) biological processes associated with Cluster 1 and performed gene set enrichment analysis (GSEA) on the pairwise comparison between COVID-19 and the combined non-COVID-19 pneumonia groups as well as between the COVID-19 and the other viral pneumonia group (**Supplemental Figure 6B, Supplemental Figure 7B, Figure 2B**, and **Figure 3B**). In both CD8^+^ and CD4^+^ T cells, genes upregulated in COVID-19 (Cluster 1) were associated with T cell proliferation, heightened immune cell activation typified by an enriched interferon (IFN)-γ and TNF-α signature, vascular-specific pathways, and co-inhibitory markers (**Supplemental Figure 6C-D**, **Supplemental Figure 7C**, and **Supplemental Files 5-10**).

**Figure 2.**
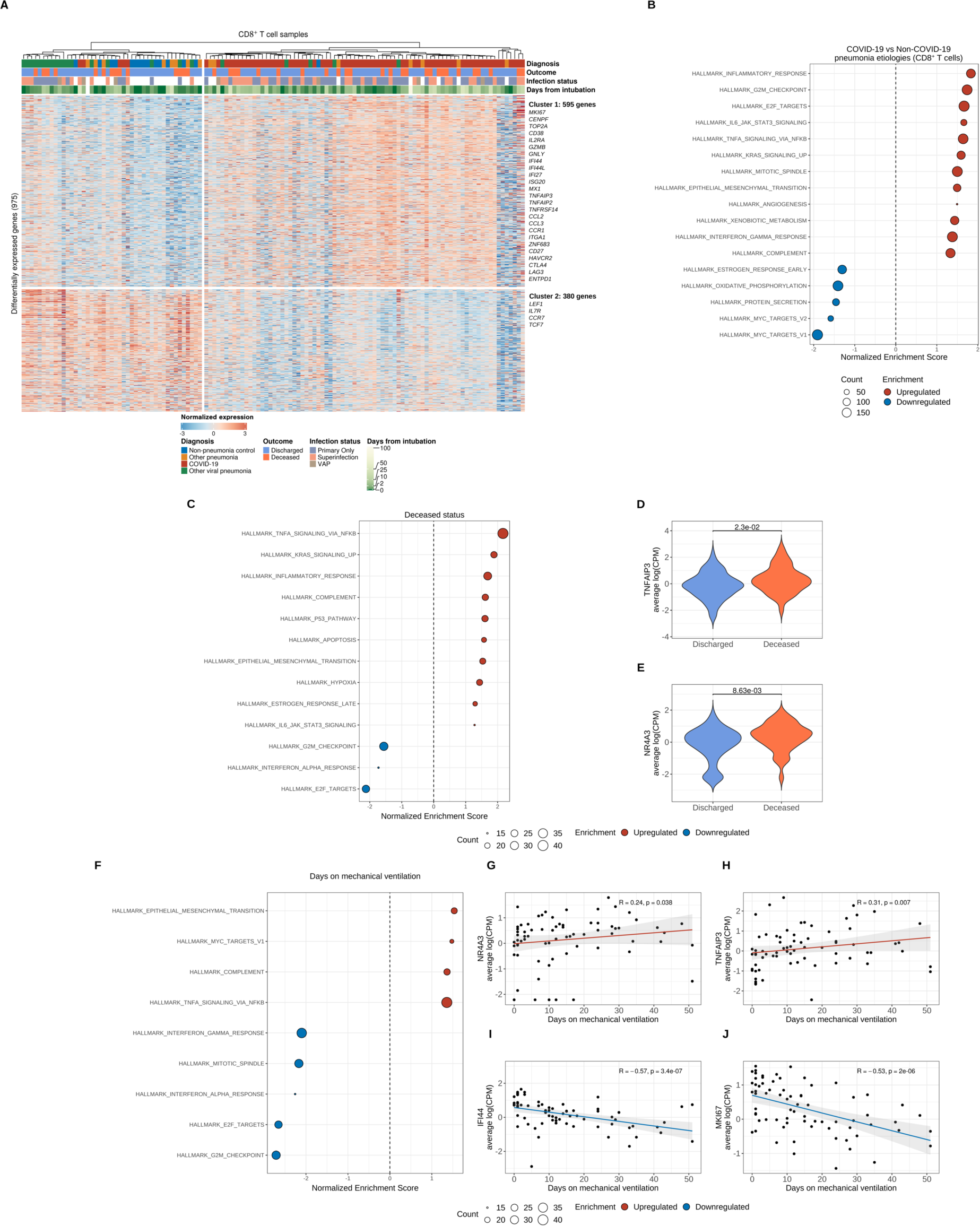
Transcriptional profiling of alveolar CD8^+^ T cells reveals distinct effector activation signatures that are associated with clinical outcomes that shift throughout the course of severe SARS-CoV-2 pneumonia. **(A)** *K*-means clustering of 975 differentially expressed genes (*q* < 0.05, likelihood-ratio test with FDR correction) across pneumonia diagnoses. Columns represent unique samples and column headers are color-coded by diagnosis, binary outcome (whether a given patient was discharged from the hospital alive or died during hospitalization), duration of mechanical ventilation (blanks indicate chronically ventilated patients), and infection status (presence or absence of bacterial superinfection in patients with COVID-19 or other viral pneumonia). The VAP (ventilator-associated pneumonia) flag designates samples from non-pneumonia controls or patients with COVID-19 or other viral pneumonia who cleared the virus and then developed a bacterial pneumonia. Samples were clustered using Ward’s minimum variance clustering method. Representative genes are shown for each cluster. **(B)** Gene set enrichment analysis (GSEA) of Hallmark gene sets for the pairwise comparison between COVID-19 samples and combined non-COVID-19 samples (non-pneumonia control, other pneumonia, and other viral pneumonia). Count denotes pathway size after removing genes not detected in the expression dataset. Enrichment denotes significant (*q* < 0.25 with FDR correction) upregulated (red) and downregulated (blue) pathways by normalized enrichment score. **(C)** GSEA of genes from COVID-19 samples after performing correlation analysis of differentially expressed genes in CD8^+^ T cells and binary outcome variable with Spearman rank correlation coefficient computation. Count denotes pathway size after removing genes that were not detected. Enrichment denotes significant (*q* < 0.25 with FDR correction) upregulated (red) and downregulated (blue) pathways by normalized enrichment score (NES). **(D-E)** Leading edge analysis reveals selected core genes driving pathway enrichment signal in the binary outcome variable. **(F)** GSEA of COVID-19 samples after performing correlation analysis of differentially expressed genes in CD8^+^ T cells and the duration of mechanical ventilation variable with Spearman rank correlation coefficient computation. Count denotes pathway size after removing genes that were not detected. Enrichment denotes significant (*q* < 0.25 with FDR correction) upregulated (red) and downregulated (blue) pathways by normalized enrichment score. **(G-J)** Leading edge analysis reveals selected core genes driving pathway enrichment signal in the duration of mechanical ventilation variable.

**Figure 3.**
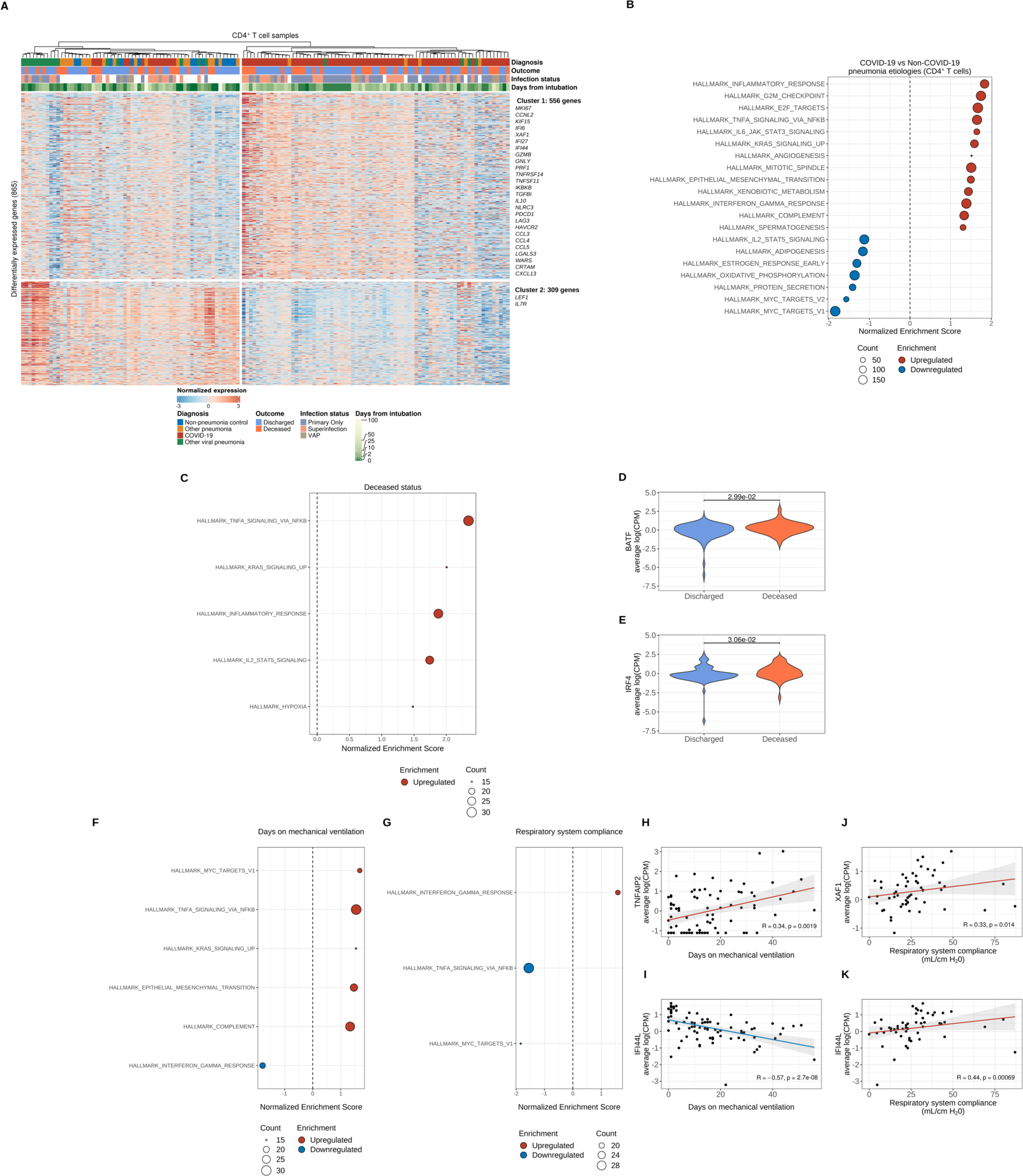
Transcriptional profiling of alveolar CD4^+^ T cells reveals distinct effector activation signatures that are associated with clinical outcomes that shift throughout the course of severe SARS-CoV-2 pneumonia. **(A)** *K*-means clustering of 865 differentially expressed genes (*q* < 0.05, likelihood-ratio test with FDR correction) across pneumonia diagnoses. Columns represent unique samples and column headers are color-coded by diagnosis, binary outcome (whether a given patient was discharged from the hospital alive or died during hospitalization), duration of mechanical ventilation (blanks indicate chronically ventilated patients), and infection status (presence or absence of bacterial superinfection in patients with COVID-19 or other viral pneumonia). The VAP (ventilator-associated pneumonia) flag designates samples from non-pneumonia controls or patients with COVID-19 or other viral pneumonia who cleared the virus and then developed a bacterial pneumonia. Samples were clustered using Ward’s minimum variance clustering method. Representative genes are shown for each cluster. **(B)** Gene set enrichment analysis (GSEA) of Hallmark gene sets for the pairwise comparison between COVID-19 samples and combined non-COVID-19 samples (non-pneumonia control, other pneumonia and other viral pneumonia). Count denotes pathway size after removing genes that were not detected. Enrichment denotes significant (*q* < 0.25 with FDR correction) upregulated (red) and downregulated (blue) pathways by normalized enrichment score. **(C)** GSEA of COVID-19 samples after performing correlation analysis of differentially expressed genes in CD4^+^ T cells and the binary outcome variable with Spearman rank correlation coefficient computation. Count denotes pathway size after removing genes not present in expression dataset. Enrichment denotes significant (*q* < 0.25 with FDR correction) upregulated (red) and downregulated (blue) pathways by normalized enrichment score. **(D-E)** Leading edge analysis reveals selected core genes driving pathway enrichment signal in the binary outcome variable. **(F-G)** GSEA of COVID-19 samples after performing correlation analysis of differentially expressed genes in CD4^+^ T cells and the duration of mechanical ventilation variable (F) and respiratory system compliance variable (G) with Spearman rank correlation coefficient computation. Count denotes pathway size after removing genes not present in expression dataset. Enrichment denotes significant (*q* < 0.25 with FDR correction) upregulated (red) and downregulated (blue) pathways by normalized enrichment score. **(H-I)** Leading edge analysis reveals selected core genes driving pathway enrichment signal in the duration of mechanical ventilation variable. **(J-K)** Leading edge analysis reveals selected core genes driving pathway enrichment signal in the respiratory system compliance variable.

Protective cellular immunity requires activation, migration, and polarization of T cells into appropriate effector programs tailored to control the immune response elicited by a given pathogen (11). In COVID-19, early induction of IFN-γ-producing T cells in the peripheral blood (type 1 immune response) has been linked to more favorable clinical outcomes, attributed to these cells’ capacity to accelerate viral clearance (24, 25). Hence, we conducted pairwise comparisons between early (≤48 hours following intubation) versus late (>48 hours following intubation) samples of alveolar CD4^+^ and CD8^+^ T cells obtained from patients with COVID-19, observing higher expression of genes involved in interferon-mediated signaling early in the course of mechanical ventilation for severe SARS-CoV-2 pneumonia (**Supplemental Figure 8A-B**). Longitudinal analysis of these interferon-stimulated genes in combined CD4^+^ and CD8^+^ T cells revealed greater expression in samples obtained early following intubation (**Supplemental Figure 8C**). Correspondingly, viral loads were higher in early compared with late samples (**Supplemental Figure 8D**). The magnitude and kinetics of viral load have been associated with disease severity and mortality (13, 26, 27). Notably, patients who succumbed to COVID-19 had a higher initial viral load with a slower decline compared with patients who survived their hospitalization (**Supplemental Figure 8D-F**). In most patients, viral load declined or the BAL fluid became PCR negative for SARS-CoV-2 over the course of COVID-19, similar to observations of sputum samples (28).

We then analyzed BAL fluid T cell transcriptomic signatures by performing correlation analyses between differentially expressed genes in CD8^+^ and CD4^+^ T cells and clinical variables followed by GSEA with leading edge analysis. We found that T cells from patients with poor outcomes from severe SARS-CoV-2 pneumonia exhibited upregulation of Hallmark processes linked to inflammatory responses and TNF-α/NF-κB signaling and downregulation of processes associated with cell proliferation and IFN-γ signaling (**Figure 2C-J**, **Figure 3C-F and H-I, Supplemental Figure 7D-G**, and **Supplemental Figure 9A-J**). Conversely, higher respiratory system compliance (a marker of normal lung function) was associated with greater expression of cell proliferation and interferon-related genes and downregulation of TNF-α signaling via NF-κB (**Figure 3G and J-K** and **Supplemental Figure 9K-O**).

CD4^+^FOXP3^+^ regulatory T (Treg) cells populate the alveolar space following virus-induced lung injury and orchestrate tissue-protective and reparative mechanisms (29). Hence, we performed *K*-means clustering of 80 differentially expressed genes in Treg cells (FDR *q* < 0.05) and identified two clusters that distinguished patients with COVID-19 from other groups (**Supplemental Figure 10A**). Notably, GSEA of the pairwise comparison between patients with COVID-19 and combined patients in the non-COVID-19 pneumonia groups revealed enrichment of processes linked to cell proliferation **Supplemental Figure 10B**). Comparing patients with COVID-19 to the other viral pneumonia group identified enrichment in interferon signaling (**Supplemental Figure 10C**). Most lung T cells are T resident memory (TRM) cells, and in mouse models of SARS-CoV infection and lung fluid samples from patients with severe SARS-CoV-2 pneumonia, TRM cells have been suggested to play a protective role (19, 30). We therefore leveraged our previously published single-cell RNA-sequencing dataset (4) to perform *in silico* cell-type deconvolution of T cell bulk RNA-sequencing analysis. We found the majority of both alveolar CD8^+^ and CD4^+^ T cells to be memory T cells across all pneumonia diagnoses (**Supplemental Figure 10D-I**). In CD8^+^ T cell samples from patients with COVID-19, the TRM population remained elevated irrespective of outcome or timing of sampling. Notably, while proliferating T cells tended to decrease in late samples overall, the cytotoxic compartment tended to increase in late samples from deceased patients with COVID-19 (**Supplemental Figure 10F**). Lastly, in CD4^+^ T cell samples from patients with COVID-19, both central memory and Treg cells were more abundant than cytotoxic and proliferating subsets (**Supplemental Figure 10I**).

### Alveolar T cell receptor profiling reveals distinct patterns in patients with COVID-19

The specificity and complexity of T cell responses is determined by the T cell receptor (TCR). Accordingly, we performed bulk TCR-sequencing on CD8^+^ and CD4^+^ alveolar T cells—the bulk of which are TRM cells as suggested in the deconvolution analysis above—to ascertain TCR repertoire signatures that shape the adaptive immune response to severe pneumonia. We used RNA from 130 alveolar T cell samples that had sufficient residual RNA (>0.5 ng) to generate TCR-sequencing libraries (see **Figure 1A** and **Supplemental Figure 1B**). These samples corresponded to 47 patients with similar gender distribution, severity of illness, and mortality between groups (**Supplemental Figure 11A-F**).

The alpha diversity with clonotype richness measurement within CD8^+^ and CD4^+^ T cells combined was similar across groups, although there were small yet statistically significant differences based on pneumonia category, timing of sample acquisition relative to intubation, and patient outcome (**Supplemental Figure 12A-C**). Similar to prior reports (31-33), we found that TCR repertoire diversity trended lower as a function of age (**Supplemental Figure 12D**). Approximately 50% of patients with severe SARS-CoV-2 pneumonia develop a secondary bacterial pneumonia either as a superinfection or a *de novo* ventilator-associated pneumonia (VAP) following viral clearance (14, 15). We hypothesized that TCR repertoire diversity would become more oligoclonal (i.e., narrow) in response to new pathogens encountered during secondary bacterial pneumonias. Indeed, whereas richness increased over the course of mechanical ventilation in the non-COVID-19 groups, we found a significant decrease in richness over time in the COVID-19 group (**Supplemental Figure 12E**), consistent with the greater risk of secondary bacterial pneumonia in patients with severe SARS-CoV-2 pneumonia relative to similarly ill patients with other causes of pneumonia and respiratory failure (14, 15). Comparing patients with COVID-19 who survived to hospital discharge with those who did not, we observed a statistically significant difference in richness only in patients who were diagnosed with a secondary bacterial infection (superinfection or VAP) during their ICU course (**Supplemental Figure 12F**). Altogether, these findings support the clinical observation that secondary bacterial infections contribute to dynamic immune responses and outcomes in patients with severe SARS-CoV-2 pneumonia (14, 15).

### Patients with severe SARS-CoV-2 pneumonia share an enriched network of T cell specificity

We used the Grouping of Lymphocyte Interactions by Paratope Hotspots 2 (GLIPH2) algorithm (34) to identify disease-relevant TCRs with predicted shared antigen specificity within the large number of the TCR sequences obtained from alveolar T cells (CD8^+^ CDR3β = 37,297 and CD4^+^ CDR3β = 64,276). We applied stringent filtering criteria (**Supplemental Figure 13A**) to obtain enriched TCR clusters with the highest probability to bind similar HLA-restricted peptides across different pneumonia categories (**Supplemental Files 11 and 12**). Our network analysis for CD8^+^ and CD4^+^ T cells revealed greater shared TCR sequence similarity in the COVID-19 group when compared with the non-COVID-19 groups (**Supplemental Figure 13C-E** and **Supplemental Figure 14A-E**). Additionally, gene usage analysis revealed a distinctive enrichment of TRBV27/TRBV12-3 and TRBV20-1/TRBV6-6/TRBV10-1 in CD8^+^ and CD4^+^ T cells, respectively (**Supplemental Figure 13H-I** and **Supplemental Figure 14I-J**); these findings are consistent with observations in peripheral blood (35). Collectively, these results suggest an enriched network of alveolar T cell specificity that is peculiar to SARS-CoV-2 pneumonia.

We next sought to uncover potential targets of alveolar CD8^+^ and CD4^+^ T cell responses to SARS-CoV-2 infection. We undertook an unsupervised reverse epitope discovery approach (36) to interrogate post-GLIPH2-enriched alveolar CDR3β sequences and identify immunodominant epitope responses in patients with COVID-19 as well as other causes of pneumonia and respiratory failure. Specifically, we leveraged the Multiplex Identification of Antigen-Specific T-Cell Receptors Assay (MIRA) dataset (37) to compare our observed CD8^+^ and CD4^+^ TCR sequences with more than 135,000 high-confidence SARS-CoV-2-specific CDR3β sequences and uncover putative shared epitopes. We annotated 42.3% (506/1,196) of GLIPH2-enriched TCRs to the MIRA MHC class I dataset and 29.0% (364/1,252) of GLIPH2-enriched TCRs to the MIRA MHC class II dataset based on identical sequence similarity (**Supplemental Files 13 and 14**). Class I/Class II datasets were comprised of 269/56 target peptide pools (deconvolved to 545/251 unique epitopes) from 15/9 unique combinatorial overlapping antigenic targets of the SARS-CoV-2 proteome, respectively (**Supplemental File 15A**). We mapped TCRs from all pneumonia categories to 32.7% (88 out of 269) of MIRA Class I and 25% (14 out of 56) of MIRA Class II peptide pools (**Supplemental Files 13 and 14)**. Notably, in patients with COVID-19, we mapped 80.6% (71/88) of MIRA Class I and 85.7% (12/14) of MIRA Class II peptide pools. Altogether, these analyses support the specificity of the alveolar T cell response to SARS-CoV-2 pneumonia compared with other causes of pneumonia and respiratory failure in our cohort.

### Antigen hierarchy distribution of alveolar T cells targeting the SARS-CoV-2 proteome is associated with clinical outcomes and age throughout the course of severe SARS-CoV-2 pneumonia

In alveolar CD8^+^ T cells, analysis of SARS-CoV-2 antigenic targets demonstrated overall dominance of ORF1ab (31.6%), Spike (S, 23.2%), and Nucleocapsid (N, 22.2%) (**Figure 4A**). Combined, N and S accounted for 48.8% of epitopes in patients with COVID-19 who were discharged from the hospital, whereas ORF1ab accounted for 43.5% of epitopes in patients who died from COVID-19 (**Figure 4B**). In patients with COVID-19, we found that within the ORF1ab polyprotein complex, the non-structural proteins NSP3 and NSP12 accounted for 35.1% and 22.0% of overall targets, respectively (**Figure 4C**). Additionally, while NSP2 was significantly enriched in patients with COVID-19 who survived to hospital discharge (FDR *q* = 6.34e-03), NSP12 was higher in patients who died, but this comparison did not reach statistical significance (FDR *q* = 1.57e-01) (**Figure 4D**). In alveolar CD4^+^ T cells, we found that 95% of TCRs recognized structural proteins (M, 52.2%; S, 25.0%; N, 17.8%) (**Supplemental Figure 14A)**, consistent with the known skewing of CD4^+^ T cell responses toward viral structural proteins (38).

**Figure 4.**
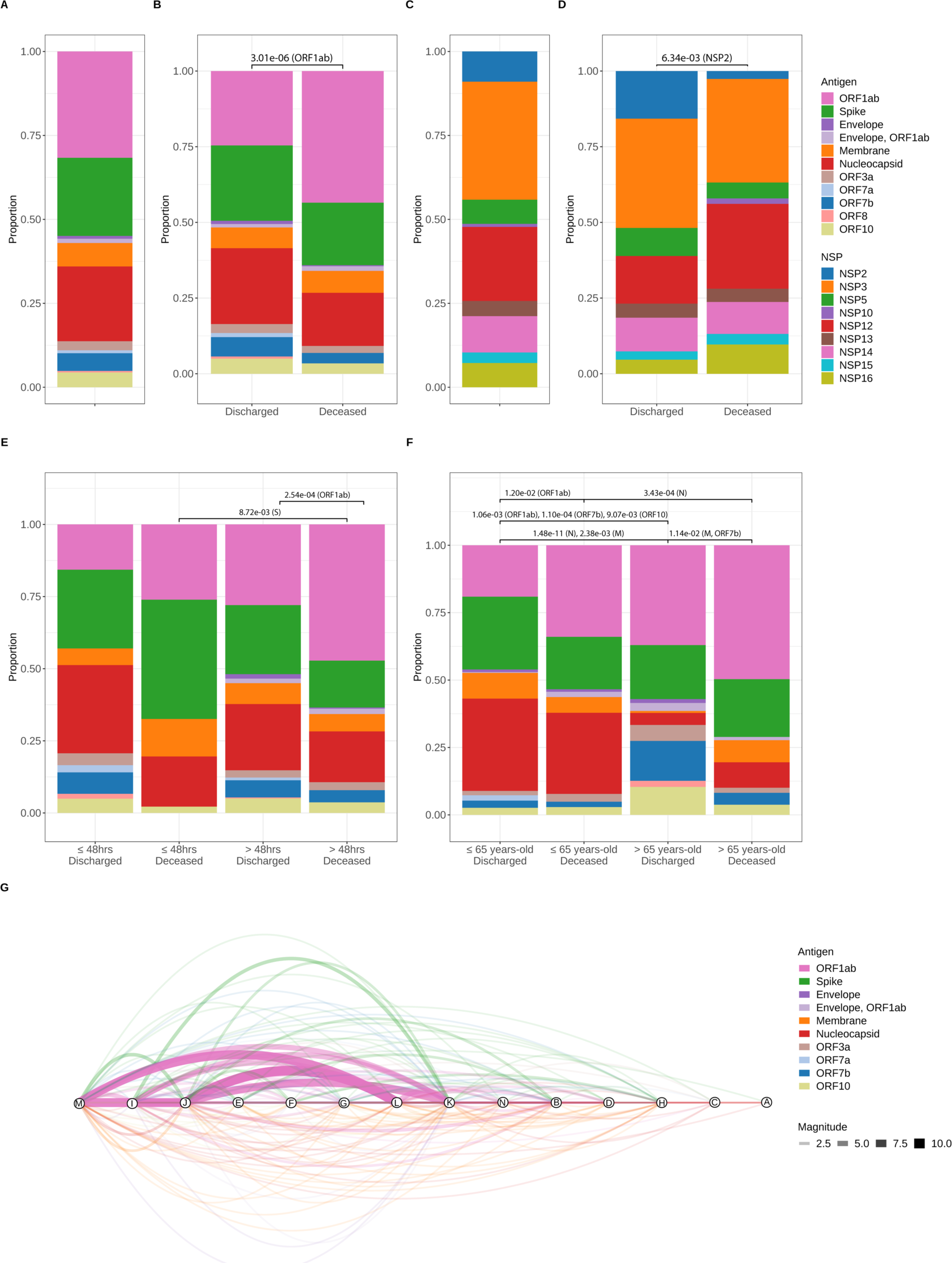
Alveolar CD8^+^ T cell targets exhibit a distinctive pattern of antigen hierarchy that is associated with clinical outcomes throughout the course of severe SARS-CoV-2 pneumonia. **(A)** Proportion of alveolar CD8^+^ T cell responses by SARS-CoV-2 protein. TCR sequences identified in samples from patients with COVID-19 were cross-referenced with the MIRA I dataset to identify reactivity against specific SARS-CoV-2 antigens. *n* of patients = 14, *n* of samples = 29. (**B**) SARS-CoV-2 antigen cross-referenced TCR sequences grouped by binary outcome. *n* of patients (Discharged = 9 and Deceased = 5), *n* of samples (Discharged = 15, Deceased = 14). *q*-value < 0.05, row wise Fisher exact tests with FDR correction (per antigen). (**C**) Nonstructural proteins (NSP) within the ORF1ab complex. *n* of patients = 13 and *n* of samples = 24. (**D**) NSP and binary outcome. *n* of patients (Discharged = 8 and Deceased = 5), *n* of samples (Discharged = 12 and Deceased = 12). *q*-value < 0.05, row wise Fisher exact tests with FDR correction (per NSP). (**E**) Timing of BAL sampling and binary outcome. *n* of patients (Discharged, ≤48 hours = 5; Deceased, ≤48 hours = 2; Discharged, >48 hours = 6; Deceased, >48 hours = 5) and *n* of samples (Discharged and ≤48 hours = 5; Deceased, ≤48 hours = 2; Discharged, >48 hours = 10; Deceased, >48 hours = 12). *q*-value < 0.05, row wise Fisher exact tests with FDR correction (per antigen). (**F**) Age and outcome. *n* of patients (Discharged, ≤65 years-old = 6; Deceased, ≤65 years-old = 2; Discharged, >65 years-old = 3; Deceased, >65 years-old = 3) and *n* of samples (Discharged, ≤65 years-old = 10; Deceased, ≤65 years-old = 5; Discharged, >65 years-old = 6; Deceased, >65 years-old = 8). *q*-value < 0.05, row wise Fisher exact tests with FDR correction (per antigen). **(G)** Network analysis of shared TCR sequences recognizing SARS-CoV-2 epitopes. Nodes represent unique patients in the COVID-19 group (labeled here A-N), edges constitute TCR sequences shared by at least 2 patients mapped to a MIRA class I dataset epitope pool, and width of edges (magnitude) denotes total number of shared TCR sequences. Edges are color-coded by SARS-CoV-2 antigens.

The antigenic evolution and breadth of alveolar T cell responses throughout the prolonged course of severe SARS-CoV-2 pneumonia remains unknown. Hence, we next sought to ascertain whether protracted recovery from illness was associated with changes in the distribution hierarchy of SARS-CoV-2 antigens. We found that in patients who recovered from COVID-19, CD8^+^ T cells maintained an S- and N-specific T cell response in both early and late samples, while patients who died had fewer S targets and exhibited ORF1ab immunodominance during the late phase of infection (**Figure 4E**). Similar to other pathogens, SARS-CoV-2 disproportionately affects older patients (39, 40). We found that samples from patients ≤65 years-old were primarily enriched for S- and N-targets, with those who died demonstrating a significantly greater proportion of ORF1ab targets (**Figure 4F**). Notably, ORF1ab was the immunodominant antigen in patients >65 years-old who died. Collectively, these results demonstrate that alveolar CD8^+^ T cells’ distinctive pattern of recognition across the SARS-CoV-2 proteome during severe SARS-CoV-2 pneumonia is associated with clinical outcome and age.

### Alveolar epitope analysis reveals distinctive patterns of immunodominance that are associated with divergent clinical outcomes in patients with severe SARS-CoV-2 pneumonia

To identify the overall breadth, immunodominance, and immunoprevalence of the epitope repertoire recognized by alveolar T cells during the course of severe SARS-CoV-2 pneumonia, we performed a network analysis uncovering the shared specificity of SARS-CoV-2 epitopes in the COVID-19 group (**Figure 4G**). Cross-referencing the MIRA MHC Class I dataset, we identified 44/71 unique peptide pools from 10/11 SARS-CoV-2 proteins that were shared in patients with COVID-19. Although the breadth of CD8^+^ T cell responses to SARS-CoV-2 antigens between discharged and deceased patients (10 discharged versus 8 deceased) and epitope pools (21 discharged versus 27 deceased) was similar, the distribution of epitope specificity to SARS-CoV-2 proteins was substantially different (**Figure 5A-F**). Specifically, we found that while immunodominant and immunoprevalent targets were well-distributed across the viral proteome in patients who recovered, targets in those who died from SARS-CoV-2 pneumonia were highly and disproportionately enriched in the ORF1ab polyprotein complex. In CD4^+^ T cells, we identified 8/12 unique peptide pools from 4/9 SARS-CoV-2 proteins shared among patients with COVID-19 **Supplemental Figure 14F-H**).

**Figure 5.**
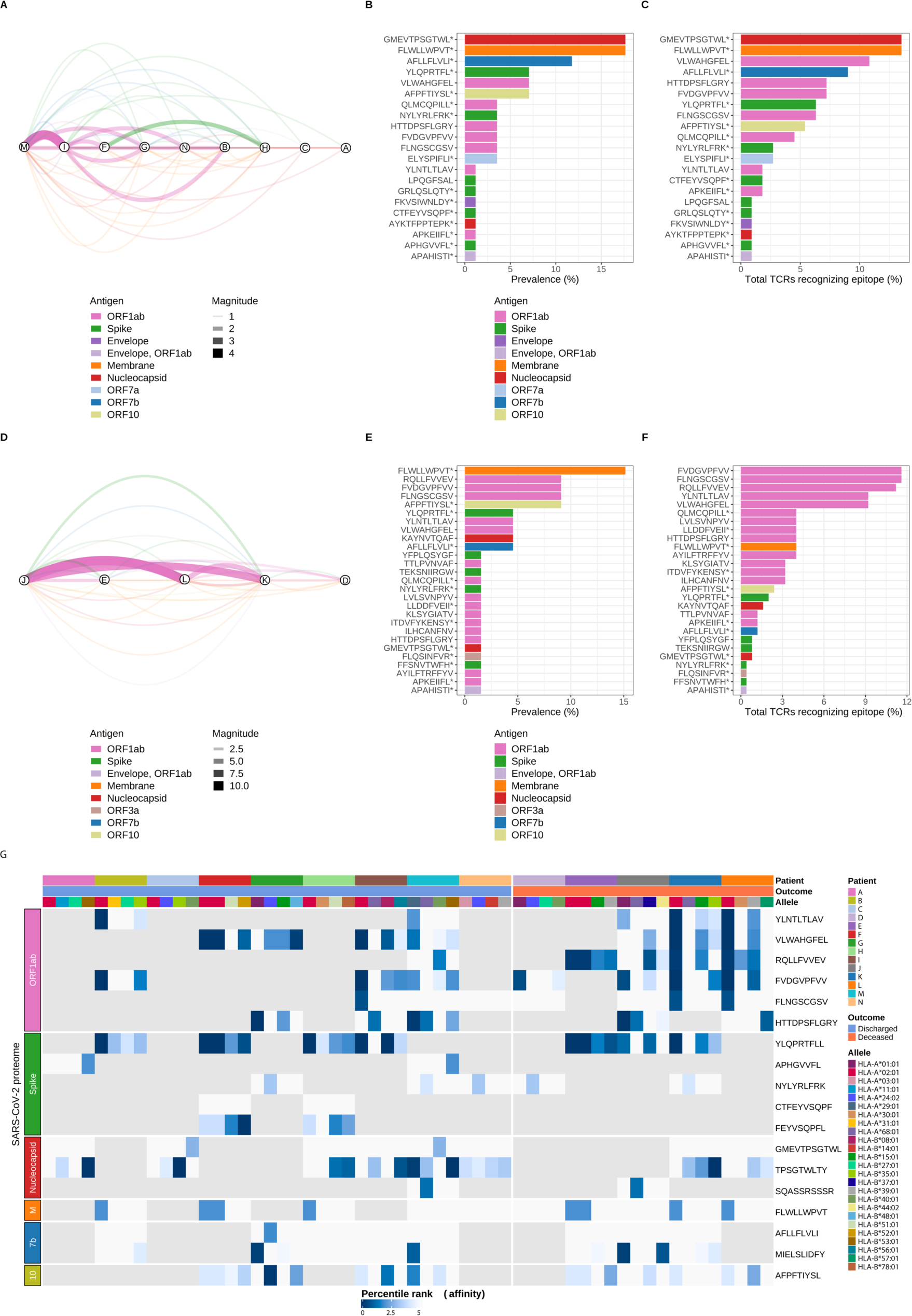
SARS-CoV-2 epitope mapping reveals a distinctive landscape of peptide immunodominance and immunoprevalence associated with outcomes in patients with COVID-19. **(A)** Network analysis of shared TCR sequences from CD8^+^ T cells recognizing SARS-CoV-2 epitopes. Nodes represent the nine unique patients with COVID-19 who survived hospital discharge (labeled here using the lettering scheme from Figure 4G), edges constitute shared TCR sequences by at least two patients mapped to a MIRA class I dataset epitope pool, and width of edges (magnitude) denotes total number of shared TCR sequences. Edges are color-coded by SARS-CoV-2 antigens. **(B)** Immunoprevalence of SARS-CoV-2 epitopes in discharged patients was calculated by counting the number of events when a given epitope was shared by at least two patients. Total counts from all 21 identified epitopes are represented as percentage (%) of TCRs recognizing a given epitope. **(C)** Overall number of TCR sequences mapped to a given SARS-CoV-2 epitope in discharged patients was calculated by counting all events of TCRs recognizing an epitope. Total counts from all 21 identified epitopes are represented as percentage (%). **(D)** Network analysis of shared TCR sequences recognizing SARS-CoV-2 epitopes in the five unique patients with COVID-19 who died during hospitalization (labeled here using the lettering scheme from Figure 4G), edges constitute shared TCR sequences by at least two patients mapped to a MIRA class I dataset epitope pool, and width of edges (magnitude) denotes total number of shared TCR sequences. Edges are color-coded by SARS-CoV-2 antigens. **(E)** Immunoprevalence of SARS-CoV-2 epitopes in deceased patients was calculated by counting the number of events when a given epitope was shared by at least two patients. Total counts from all 27 identified epitopes are represented as percentage (%). **(F)** Overall number of TCR sequences mapped to a given SARS-CoV-2 epitope in deceased patients was calculated by counting all events of TCRs recognizing an epitope. Total counts from all 27 identified epitopes are represented as percentage (%). **(G)** Heatmap of predicted SARS-CoV-2 epitope binding affinity to patient-specific HLA molecules grouped by unique patient, binary outcome, HLA alleles, and SARS-CoV-2 antigens. Percentile rank denotes predicted affinity strength with percentile ranks <1% and <5% denote strong and weak MHC binder sequences, respectively. Gray tiles represent epitopes not detected within a given patient. Column labels are color-coded by patient, binary outcome, and HLA alleles. Row labels are color-coded by SARS-CoV-2 antigens. M (Membrane), 7b (ORF7b), 10 (OFR10), * denotes other epitopes are present within MIRA class I dataset peptide pool.

TCRs recognize cognate peptides in an HLA-restricted manner; thus, HLA gene polymorphisms strongly affect population-based T cell effector responses to different epitopes (41). We used a bioinformatics tool (arcasHLA) to infer HLA typing in our COVID-19 group (**Supplemental Figure 13B and F-G** and **Supplemental Figure 14B**) (42). To predict the specificity of T cell responses, we used a machine learning tool (NetMHCIpan-4.1) (43) to assess an epitope’s likelihood of binding a given HLA molecule. We selected the top five epitopes from our peptide network analysis by binary outcome (discharged or deceased) (**Figure 5B-C and E-F**) and predicted immunodominant epitopes within the immunodominant SARS-CoV-2 region at the level of individual patients (18 epitopes total) (**Supplemental File 15B**). We then mapped predicted CD8^+^ T cell reactivities from all 14 patients in the COVID-19 group to the four inferred patient-specific HLA A and B alleles (**Figure 5G**) (44). The majority of immunodominant epitopes (17/18 epitopes, 94.4%) were predicted to bind (defined as a percentile rank < 5%) to at least one HLA molecule expressed in the corresponding patient. All ORF1ab epitopes along with epitopes YLQ (Spike), TPS (Nucleocapsid), and MIE (ORF7b) were predicted to bind with high-affinity (defined as percentile rank < 1%) to at least three patient-specific HLA molecules. Additionally, the most frequently identified dominant epitopes were predicted to bind with high affinity to multiple HLA molecules, which tend to be both immunodominant and immunoprevalent in human populations (45). Altogether, predictive mapping of SARS-CoV-2 epitopes targeted by alveolar T cells identified distinctive patterns of regional viral protein and peptide dominance, which were accurately restricted by patient-specific HLA allele expression and associated with outcomes in our cohort.

### Alveolar T cells in patients with severe SARS-CoV-2 pneumonia are predicted to recognize epitopes shared with other human coronaviruses

Previous encounters with extant human coronaviruses (HCoV), which exhibit conserved sequence homology with SARS-CoV-2, have been suggested to induce functional cross-reactive responses in patients with COVID-19 (46-53). This phenomenon is particularly relevant in older individuals in whom memory T cell subsets become relatively abundant as the numbers and diversity of the naïve T cell compartment decline with age (31-33, 54, 55). Accordingly, we selected immunodominant SARS-CoV-2 epitopes from our peptide network analysis (see **Figure 5A-F)** to calculate pairwise sequence similarity with peptides from HCoVs (OC43, HKU1, NL63, and 229E) (**Figure 6A**). Notably, the average conservation for epitopes across distinct coronavirus antigens was highest for ORF1ab when compared to Spike and Nucleocapsid antigens (**Figure 6B** and **Supplemental File 15C**). Next, we compared the frequencies of all 88 SARS-CoV-2 epitopes detected in the MIRA I dataset between patients with COVID-19 and patients with non-COVID-19-related pneumonia and respiratory failure. While some of the most immunodominant epitopes in patients with COVID-19 were also detected in unexposed individuals, the overall prevalence of these epitopes was significantly enriched in the COVID-19 group (**Figure 6C-D** and **Supplemental Figure 14K-L**). Altogether, our results suggested T cell cross-reactivity with HCoV in patients with severe SARS-CoV-2 pneumonia.

**Figure 6.**
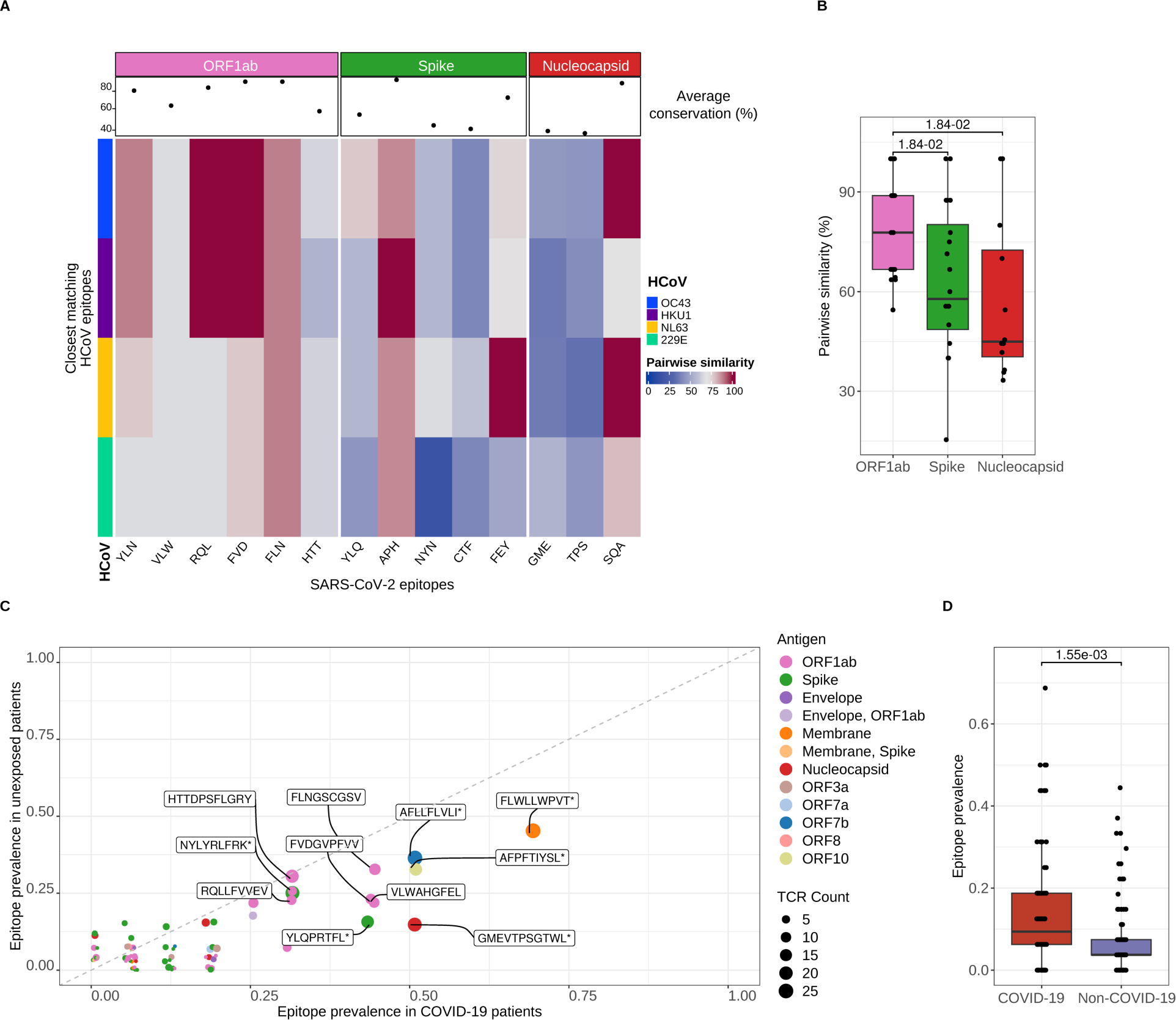
Predicted alveolar CD8^+^ T cell receptor targets cross-react with other human coronaviruses. **(A)** Heatmap of conserved sequence similarity between dominant SARS-CoV-2 epitopes detected in alveolar CD8^+^ T cells and human coronaviruses (HCoV). Columns represent SARS-CoV-2 epitopes grouped and color-coded by antigen region. Rows are color-coded by distinct HCoV. Pairwise similarity denotes percentage of sequence homology between viruses. An average sequence homology percentage across all HCoV for each SARS-CoV-2 epitope is depicted as a dot in the column header. **(B)** Pairwise sequence similarity scores between SARS-CoV-2 epitopes and closest matching epitopes from human coronaviruses. *q* < 0.05, pairwise Wilcoxon rank-sum tests with FDR correction. **(C)** Scatter plot of SARS-CoV-2 epitope prevalence of CD8^+^ T cells in patients with COVID-19 (*n* = 14) and without COVID-19 (unexposed, *n* = non-pneumonia control [4], other pneumonia [7], and other viral pneumonia [8]). Dots are color coded by SARS-CoV-2 antigen. Dot size corresponds to the number of detected TCR sequences recognizing a given antigen. Random variation to location of points was added with *geom_jitter* function from ggplot2 v.3.4.4 for improved visualization. **(D)** SARS-CoV-2 epitope prevalence in overall patient cohort from bulk CD8^+^ TCR sequencing grouped by COVID-19 status (*n* of patients = non-pneumonia control [4], other pneumonia [7], COVID-19 [14] and other viral pneumonia [8]). Wilcoxon rank sum test, *p* < 0.05.

## Discussion

Successful pulmonary immune responses to acute infection coordinate innate and adaptive immune events that lead to host recovery following severe pneumonia (56). In this study, we performed serial sampling of the alveolar spaces of patients with severe pneumonia to test the null hypothesis that dynamic pulmonary T cell responses are similar between causes of pneumonia (57). Rejecting the null hypothesis, we found that expansion of monocytes and T cells exhibiting enriched interferon signaling pathways within the alveolar spaces of patients with severe SARS-CoV-2 pneumonia early in the course of mechanical ventilation correlated with survival to hospital discharge, a pattern that was not observed in patients with pneumonia and respiratory failure due to other causes. In contrast, higher levels of persistent T cell activation with a TNF-α/NF-κB inflammatory signature were associated with poor outcomes, including mortality. TCR repertoire profiling of sorted CD8^+^ and CD4^+^ T cells revealed distinct specificity networks in patients with COVID-19 and identified patterns of immunodominant SARS-CoV-2 antigenic regions at the level of specific epitopes and HLA alleles that were associated with clinical outcome, age, and cross-reactivity with other human coronaviruses. Interestingly, we found that ORF1ab antigen specificity was associated with hospital mortality and age in patients with severe SARS-CoV-2 pneumonia. Altogether, these data identify unique T cell responses that evolve over time in the lungs of mechanically ventilated patients with COVID-19 that are distinct from other causes of pneumonia in similarly ill patients.

Preclinical and human studies have investigated how dynamic changes in T cell-specific activation programs determine Sarbecovirus disease severity (30, 58-60). While an early and robust interferon response in T cells correlates with effective viral clearance and mild disease, a delayed and suboptimal interferon response is associated with impaired T cell function, leading to vascular permeability and the induction of an unchecked innate immune-mediated proinflammatory response (22, 25, 58, 61, 62). Our unique cohort allowed us to identify the transcriptional response of alveolar T cells in unvaccinated patients with COVID-19 over the course of their disease and compare them to similarly ill patients with pneumonia attributable to other pathogens. We found that an interferon response-dominated T cell molecular signature within the first 48 hours following intubation was enriched in patients with COVID-19 who went on to experience favorable clinical outcomes, including survival. Conversely, progressive SARS-CoV-2 pneumonia in patients with persistent respiratory failure was characterized by a decline in interferon responses and dominated by proinflammatory T cell responses driven by signaling through TNF-α and NF-κB. Clinically, patients with persistent respiratory failure and NF-κB activation had higher disease severity and risk for secondary bacterial pneumonias. T cells from these patients expressed inhibitory molecules (e.g., *PDCD1*, *HAVCR2*, *LAG3*) and transcription factors (*NR4A3*, *IRF4*, *PRDM1*) that have been linked to T cell exhaustion but in this context are more likely indicative of persistent activation (63). Collectively, these results suggest that failure of interferon responses early in the course of critical illness leads to persistent activation of NF-κB-driven inflammation over the course of prolonged respiratory failure, perhaps associated with acquisition of secondary bacterial pneumonia.

An open question in the field is how SARS-CoV-2 proteome-specific T cell targets evolve throughout the course of COVID-19 and whether any of these targeted antigenic regions are associated with disease outcome (64). Our TCR analysis demonstrated that while early CD8^+^ T cell samples from individuals ≤65 years-old with favorable outcomes exhibited a higher proportion of targets against structural proteins (Spike and Nucleocapsid), late samples from patients >65 years-old who died from SARS-CoV-2 pneumonia were highly enriched for the non-structural proteins within the ORF1ab polyprotein complex. We were able to further narrow the predicted binding affinity of T cells for specific SARS-CoV-2-immunodominant epitopes in an HLA-restricted fashion and predict cross reactivity with other endemic human coronaviruses. We posit that ORF1ab-specific T cells during the late phase of severe SARS-CoV-2 pneumonia could be associated with augmented immune escape mechanisms leading to delayed adaptive T cell responses in a subset of patients exhibiting poor outcomes.

Multiple studies have shown that blood T cells collected from individuals before the COVID-19 pandemic possess immune reactivity against specific SARS-CoV-2 peptides in regions of conserved homology with human coronaviruses, including ORF1ab, the S2 region of Spike, and Nucleocapsid (52, 65-67). These observations led to the hypothesis that prior encounters with human coronaviruses—particularly in children and younger adults—could mediate protective immune responses. Nevertheless, the functional characteristics of cross-reactive immunity and their impact on COVID-19 clinical outcomes remain unclear. Bacher and colleagues demonstrated that in previously SARS-CoV-2-unexposed individuals with COVID-19, pre-existing SARS-CoV-2-reactive circulating T cells possessed low TCR avidity and were enriched in patients with more severe disease, arguing against a protective function for cross-reative T cells (49). Furthermore, cross-reactivity might not only be limited to human coronaviruses (46, 49), and others have shown that microbial peptides from commensal bacteria and other viruses are potential sources of heterologous immunity to SARS-CoV-2 (68, 69). Here, we predicted that TCRs specific for immunodominant SARS-CoV-2 peptides in our dataset cross-react with other human coronaviruses. Hence, while some level of cross-reactivity may be protective, it is tempting to speculate that in certain patients with severe COVID-19, low-avidity, cross-reactive T cells generated following exposure to either human coronaviruses or other microbial peptides in the lung drive dysfunctional adaptive T cell immunity. Future studies are needed to assess the functional characteristics of cross-reactive T cell responses during severe SARS-CoV-2 pneumonia.

Based on direct observations of cells obtained from the alveolar space, the data presented here support the previously proposed model of SARS-CoV-2 pathogenesis in which activated CoV-reactive T cells drive feed-forward circuits with alveolar macrophages to sustain alveolar inflammation, respiratory failure, and risk for secondary bacterial pneumonia and other ICU complications in mechanically ventilated patients with COVID-19 (4-8, 14, 15). While early expansion of T cells enriched for interferon signaling pathways that target structural SARS-CoV-2 proteins (Spike and Nucleocapsid) were associated with favorable clinical outcomes, late T cell activation was associated with expression of TNF-α/NF-κB signaling pathways, specificity directed toward the non-structural ORF1ab polyprotein complex, and mortality. Identifying these distinctive features of severe SARS-CoV-2 pneumonia may inform the design of clinical trials of agents and vaccine targets demonstrated to be effective in COVID-19 for patients with other causes of pneumonia.

## Limitations

Our study has important limitations. First, while we were able to perform deep molecular phenotyping of T cell subsets, cell numbers and study design limited our ability to perform functional studies of T cell avidity and cross-reactivity. Second, as pneumonia is an encompassing syndrome, our single-site study population was heterogeneous. We minimized this heterogeneity by focusing on an ICU population and performing careful clinical phenotyping of patients by pneumonia category, superinfection status, and clinical endpoints. Third, molecular phenotyping in the cohort was feasible only for samples with sufficient numbers of T cells present in BAL fluid. This limitation may have imparted a degree of selection bias to the analysis, although the clinical features of the patients whose samples had sufficient material for molecular phenotyping resembled the larger cohort. Fourth, because all samples were obtained early in the COVID-19 pandemic—ahead of vaccination, prior exposure, and the emergence of viral variants of concern—we cannot determine the effects of these variables on T cell responses in our cohort. At present, the majority of patients with severe SARS-CoV-2 pneumonia remain unvaccinated (70), similar to the patients in our study. Nevertheless, to the best of our knowledge, all SARS-CoV-2 infections in our cohort were primary infections rather than recurrent or breakthrough infections, which may be more common presently. Finally, our TCR analysis relied on reverse epitope prediction, which is a powerful technique to identify the likely antigens recognized by the TCRs in our dataset but is nonetheless reliant on the robustness and accuracy of existing databases. Accordingly, we have made our data publicly available for reanalysis as these techniques evolve.

## Methods

### Human participants

The details of participant recruitment in the SCRIPT Systems Biology Center have been previously reported (4, 14, 15). In brief, the SCRIPT study screened patients at least 18 years of age who were receiving mechanical ventilation and had clinical suspicion of pneumonia based on clinical signs, including, but not limited to, fever, radiographic infiltrate, and respiratory secretions and had undergone at least one BAL procedure to evaluate the presence and microbial etiology of pneumonia. For this study, we included data and samples from patients enrolled in SCRIPT from June 2018 to August 2020 in the ICU at Northwestern Memorial Hospital in Chicago. We selected this time period because it was a study era in which SCRIPT used flow cytometry to analyze and sort T cell subsets for bulk transcriptional profiling. Participants could re-enter the study under a new study identifier if they were discharged from the hospital and subsequently re-admitted to the ICU. The etiology of pneumonia (SARS-CoV-2, other viral pneumonia, other pneumonia, or non-pneumonia [intubated for reasons other than pneumonia]) and outcome of each pneumonia episode (cured, indeterminate, or not cured) were adjudicated by consensus of pulmonary and critical care medicine physicians using a validated procedure (15, 21). The etiology of pneumonia was determined based on clinical and BAL fluid data obtained on the date of study enrollment. Detailed definitions of pneumonia episodes and resolution status are available in reference (21). In this study, patients who underwent lung transplantation for refractory respiratory failure and patients who were transferred to home or inpatient hospice were adjudicated as having died in the reported binary outcome (discharged from the hospital alive versus deceased). Superinfection was defined as bacterial infection co-occurring with a viral pathogen diagnosed by BAL sampling. For the purposes of this report, ventilator-associated pneumonia (VAP) refers to incident bacterial pneumonia occurring after at least 48 hours of mechanical ventilation in patients who were enrolled as a non-pneumonia control or who were enrolled with viral pneumonia (due to SARS-CoV-2 or another virus) but who did not have evidence of an underlying viral infection at the time of the current BAL sample (i.e., had cleared their viral infection). Demographics, clinical data, and outcomes were extracted from the electronic health record (EHR) via the Northwestern Medicine Enterprise Data Warehouse (71). Racial groups with fewer than five individuals were censored to ‘Other’ to protect patient anonymity. Comorbidities were extracted based on ICD codes as aligned to Charlson Comorbidity Index at time of hospital admission. The Northwestern University Institutional Review Board approved all research involving human participants under study STU00204868. All study participants or their surrogates provided informed consent.

### Clinical management

Bronchoscopic or non-bronchoscopic BAL sampling was performed using standard techniques with modifications to limit the generation of infectious aerosols (72). We routinely instill 120 mL of non-bacteriostatic saline in four 30-mL aliquots, discarding the return from the first aliquot. Quantitative bacterial cultures, multiplex or targeted PCR (BioFire® FilmArray® Pneumonia (PN) Panel, targeted SARS-CoV-2 PCR, and Respiratory Pathogen Panel), and automated cell count and differential were performed on BAL fluid, and nearly all patients underwent urinary antigen testing for *Streptococcus pneumoniae* and *Legionella pneumophilia* serogroup 1. Patient management was guided by institutional practice, including adherence to lung-protective mechanical ventilation strategies and use of prone positioning and ECMO, consistent with published guidelines (73-77). Some patients were enrolled in the ACTT-1 placebo-controlled trial of remdesivir for COVID-19 (78) as noted in Supplemental Table 1.

### Flow cytometry analysis and sorting

The details of the standard operating procedures of the SCRIPT study for flow cytometry have been previously reported (4). In brief, BAL fluid samples were stored at 4 °C for no longer than 24 hours before filtration through a 70-µm filter, centrifugation, and hypotonic lysis (BD PharmLyse). All cell counts were performed on a K2 Cellometer (Nexcelom) with AO/PI reagent. Fc receptors were blocked using Human TruStain FcX (Biolegend) in MACS buffer (Miltenyi Biotech). Cells were incubated with fluorochrome-conjugated antibodies at 4 °C for 30 minutes, washed, and resuspended in MACS buffer containing SYTOX Green viability dye (ThermoFisher). A FACS Aria III SORP with 100-µm nozzle operating at 20 psi was used to sort pre-defined T cell populations into 300 µL of MACS buffer using previously validated gating strategies (72). Specifically, we defined regulatory T (Treg) cells as live FSC^lo^SSC^lo^ CD3ε^+^CD4^+^CD25^hi^CD127^lo^ cells, CD4^+^ non-Treg cells as live FSC^lo^SSC^lo^ CD3ε^+^CD4^+^ cells not in the Treg cell gate, bulk CD4^+^ T cells as live FSC^lo^SSC^lo^ CD3ε^+^CD4^+^ cells, and CD8^+^ T cells as live FSC^lo^SSC^lo^ CD3ε^+^CD8^+^ cells. Sorting for Treg cells, CD4^+^ non-Treg cells, and CD8^+^ T cells was performed if at least 1,000 Treg cells could be captured; otherwise, the gate was collapsed to capture bulk CD4^+^ T cells. Lastly, samples were not processed for flow cytometry if <4 mL were available for research use, if staff were unavailable, if the sample was grossly purulent (pus), or if the sample was collected in the first weeks of the pandemic before biosafety approval for research on these samples was obtained.

### Bulk RNA-sequencing and processing

Following sorting of T cell subsets, cells were pelleted by centrifugation and lysed in 350 µL of RLT Plus lysis buffer (Qiagen) supplemented with 1% 2-mercaptoethanol and immediately stored at -80 °C. The Qiagen AllPrep DNA/RNA Micro Kit was used for simultaneous isolation of RNA and DNA, and RNA quality and quantity was assessed using a 4200 TapeStation System (Agilent Technologies). RNA-sequencing libraries were prepared from 300 pg of total RNA using SMARTer Stranded Total RNA-seq Kit v2 – Pico Input Mammalian (TakaraBio). Libraries were pooled and sequenced on a NextSeq 500 instrument (Illumina), 75 cycles, single-end, to an average sequencing depth of 20.83 million reads. Computational and bioinformatics pipelines were performed using Northwestern University’s Quest High Performance Computing Cluster Facility. The pipelines were constructed based on open-source software using nf-core /rnaseq pipeline v.3.3 implemented in Nextflow v.21.04.3 (79, 80). Nf-core/rnaseq pipeline was run with nu_genomics profile and skip_bigwig option and otherwise default options. Briefly, Quality control using FastQC v.0.11.9 and adaptor trimming using Trimgalore v.0.6.6 were performed on sequence reads from the RNA-sequencing data. STAR v2.6.10d (81) was used to align the reads to the reference genome (GRCh38 version of iGenomes reference, originally donwloaded from NCBI Homo sapiens Annotation Release 106), and Salmon v.1.4.0 (82) was used for gene and transcripts quantification. Downstream analysis was performed in R v.4.1.1. Sample swaps and mis-annotations were first identified by comparing a given patient’s known sex with sex determined by levels of *XIST* and *RPSY41*, followed by exploration of expression of canonical markers for T cells subsets and macrophages (*CD8A, CD4, FOXP3, C1QC*). Samples exhibiting either poor alignment, unexpected correlation, or extreme deviation in PCA were excluded from downstream analysis. Details for these procedures and all code described below are available in our GitHub repository: https://github.com/NUPulmonary/2023_Tcell_responses.

### Differential expression analysis (DEA)

DEA was performed using edgeR v.3.36.0 (83). In brief, genes with very low count reads were filtered executing the function *filterByExpr*. Effective or normalized library sizes were calculated using *calcNormFactors* function. A generalized linear model (GLM) framework was used for matrix design with pneumonia category as imputed factor for **Figures 2A, 3A and Supplemental Figures 6A, 7A** and timing of BAL relative to intubation (COVID-19 samples only) as imputed factor for **Supplemental Figures 8A-B**. Common and tagwise dispersion were calculated using the *estimateDisp* command, and DEA was performed using likelihood ratio tests with functions *glmFit* and *glmLRT*. Significantly variable genes (FDR *q-*value < 0.05) were identified after performing a likelihood ratio test under the GLM framework as described above. We used k-means algorithms to cluster the significantly variable genes with the optimal number of clusters calculated using *fviz_nbclust* function from factoextra v.1.0.7 followed by clustering with Hartigan-Wong algorithm from *kmeans* function in stats v.4.1.1. Samples were subsequently hierarchically clustered by using Euclidean distance and Ward’s linkage method and visualized with ComplexHeatmap v.2.10.0 (84).

### Functional enrichment analysis

Gene Ontology (GO) Enrichment Analysis was performed with topGO v.2.46.0 (85). DEG were used as “genes of interest” within the gene universe. Gene Ontology annotations and attributes of interest were extracted with biomaRt v.2.50.3 (86) to build the topGO data object. Significant GO terms were obtained after performing a classical enrichment analysis (algorithm = *“classic”*, statistical test = “*Fisher exact test”*) followed by multiple testing correction using the Benjamini-Hochberg method (see **Supplemental Data Files 5-6 and 9** for all significant GO terms). To minimize redundancy between identified GO terms, help infer biological significance, and improve visualization of multiple significant GO terms, we used rrvgo v.1.6.0 (87) to group similar GO terms by semantic similarity methods (method = *Rel*, threshold = 0.5) (see **Supplemental Data Files 7-8 and 10** with all GO terms and parent terms). Gene set enrichment analysis (GSEA) (88) was performed after retrieving the Hallmark gene set collection (h.all.v7.5.1.symbols.gmt) and using log-fold change ranked genes with *fgseaMultilevel* feature, which implements an adaptive multilevel splitting Monte Carlo approach for enhanced estimation of small p-values with Fast Gene Set Enrichment Analysis (fgsea) v.1.20.0.26771021 (89).

### Correlation analysis of bulk RNA sequencing from T cell subsets with clinical outcomes

DEG (975 genes for CD8+ T cells and 865 for CD4+ T cells) obtained from DEA with edgeR were used to calculate correlation between gene expression levels and clinical metadata in COVID-19 samples. *Cor* function from stats v.4.1.4 was used to compute correlation coefficient in the presence of missing values (method = *“spearman”*, use = *“pairwise.complete.obs”)*. Gene-associated correlation coefficients were ranked for individual clinical variables of interest, followed by GSEA as described above. Genes of interest were selected from the leading edge subsets within Hallmark processes.

### Real Time SARS-CoV-2 PCR Ct values

Viral RNA was extracted from specimens using the QIAamp Viral RNA Minikit and the QIAamp 96 Virus QIAcube HT Kit (Qiagen). Viral transport media (VTM)-only controls were included in each extraction. Laboratory testing for the presence of SARS-CoV-2 was performed by quantitative reverse transcription and PCR (qRT-PCR) with the CDC 2019-nCoV RT-PCR Diagnostic Panel utilizing N1 and RNase P probes as previously described (https://www.cdc.gov/coronavirus/2019-ncov/lab/rt-pcr-panel-primer-probes.html).

Positive and negative controls for SARS-CoV-2 and Rnase P were included in each qRT-PCR experiment alongside the VTM only sample from the RNA extraction, a no template control, and standard curves for SARS-CoV-2 and Rnase P. Specimens with Rnase P cycle thresholds (Ct) above 35 were of insufficient quality and were excluded from future studies. N1 Ct values less than or equal to 35 were considered positive, and these Ct values were used in all subsequent analyses.

### Cell-type deconvolution of bulk RNA sequencing T cell signatures

Deconvolution of T cell bulk RNA sequencing was performed using AutoGeneS v.1.0.4 (90). Signatures were derived using the single cell dataset from Grant et al. *Nature.* 2021 (4). A model was trained on CD8^+^ T cells, CD4^+^ T cells, and Treg cells. Signatures were automatically derived from 1,000 highly variable genes with function *optimize* (ngen=2000, seed=0, nfeatures=200, mode=“fixed”) for CD8^+^ T cells and function *optimize* (ngen=2000, seed=0, nfeatures=150, mode=“fixed”) for CD4^+^ T cells and Treg cells. The model was then applied to bulk RNA-sequencing data to estimate the proportion of specific cell types using regression analysis.

### T cell receptor (TCR) sequencing and analysis

We performed TCR-sequencing on selected samples of CD4^+^ and CD8^+^ T cells that contained at least 0.5 ng of residual RNA following bulk RNA-sequencing. RNA quality and quantity were measured using a 4200 TapeStation using high-sensitivity RNA ScreenTape (Agilent Technologies) before library preparation using the SMARTer Human TCR α/β Profiling Kit (Takara Bio). In brief, this kit uses a 5’ RACE-like approach to capture complete V(D)J variable regions of TCR transcripts and primers that incorporate Illumina-specific adaptor sequences during cDNA amplification. Libraries were then pooled, denatured, and diluted to a 1.8-pM DNA solution. PhiX control was spiked in at 20%. Libraries were sequenced on an Illumina NextSeq 500 instrument using NextSeq 500/550 Mid Output Kit v2.5 (300 cycles) with a target read depth of approximately 17.39 million aligned reads per sample. Raw sequencing reads in FASTQ format were aligned against a default reference database of V-, D-, J- and C-gene segments, followed by assemblage into clonotypes using MiXCR software v.3.0.13 (91). MiXCR-processed files were exported and analyzed using the R-based package, immunarch v.0.9.0 (92). Clonotype diversity was calculated with the *repDiversity* function, using a nonparametric asymptotic estimator method of species richness *chao1*.

### HLA typing from RNA-sequencing

We used arcasHLA v.0.4.0 to perform high-resolution HLA class I and class II genotyping from RNA sequencing (42). Specifically, HLA sequences were extracted from mapped chromosome 6 reads in sorted BAM files (RNA sequencing files) and referenced with IMG/HLA database v.3.47.0. Singularity container v.3.8.7 was used to perform analysis using commands *arcasHLA extract* and *arcasHLA genotype*. In certain cases, the specific HLA protein field was reduced to the most frequent within the allele group.

### Identification of TCR motifs and shared specificity groups using GLIPH2

The GLIPH2 algorithm was implemented to identify TCR sequences predicted to bind similar epitopes in an HLA-restricted manner (34). After imputing participant-specific CDR3β amino acid sequences, TRβV genes, and HLA alleles from our cohort, the GLIPH2 algorithm compared CDR3β sequences to a reference database of over 200,000 nonredundant naïve CD4^+^ and CD8^+^ TCRβ sequences from 12 healthy controls and clustered them into specificity groups (patterns) according to global and local convergence sequence metrics. To identify high-confidence TCR specificity groups shared among our cohort, GLIPH2 analysis provided statistical measurements that identify TRβV gene usage bias and HLA allele usage, comparing enriched TCR sequences between our dataset and a reference dataset using Fisher exact test. Accordingly, we used these variables coupled with analysis of clusters containing at least 3 unique CDR3β sequences, from at least 3 distinct patients within our cohort (**Supplemental Figure 13A**).

### TCR network analysis and COVID-19-specific epitope dominance analysis

To identify high-confidence SARS-CoV-2-specific TCR epitopes, we mapped our cohort’s CDR3β sequences to the MIRA dataset (37). To visualize shared TCR specificity between pneumonia diagnoses, we used network v.1.18.1, tidygraph v.2.3, and ggraph v.2.1.0 (93-95).

### Selection of SARS-CoV-2 dominant epitopes and sequence conservation analysis

For **Figure 5G**, we selected both the top five SARS-CoV-2-specific epitopes from our peptide network analysis (**Figure 5B-C and E-F**). We also selected immunodominant epitope(s) within the immunodominant SARS-CoV-2 antigenic region(s) at the level of individual patients (18 epitopes total, **Supplemental File 15B**), followed by implementation of NetMHCpan EL 4.1 tool (available on Immune Epitope Database and Analysis Resource, http://tools.iedb.org/mhci/) (43) to predict their potential binding affinity in a patient-specific, HLA-restricted manner. For epitopes with poor binding capacity (e.g., CTFEYVSQPF, GMEVTPSGTWL and AFLLFLVLI), alternate epitopes within their respective MIRA class I epitope pools were used for analysis (e.g., FEYVSQPFL, TPSGTWLTY and MIELSLIDFY, respectively). To estimate the sequence conservation between SARS-CoV-2 epitopes and other HCoV-related epitopes, we first obtained whole-genome sequences for SARS-CoV-2 (GenBank ID: MN985325.1), HCoV 229E (GenBank ID: MN306046.1), HCoV HKU1 (GenBank ID: KY983584.1), HCoV NL63 (GenBank ID: KX179500.1), and HCoV OC43 (GenBank ID: MN306053.1) from the NCBI database. Next, we calculated the pairwise sequence similarity score (https://www.ebi.ac.uk/Tools/psa/) for each selected SARS-CoV-2 epitope. Specifically, we imputed the sequence of a given selected SARS-CoV-2 epitope against the whole antigen-specific and matching HCoV sequence by using global alignment and following these parameters: EMBOSS Needle, Needleman-Wunsch algorithm, BLOSUM62 matrix, gap open (10), gap extend (0.5), end gap penalty (false), end gap open penalty (10) and end gap extension penalty (0.5). The resultant matching HCoV epitopes were subsequently imputed to calculate the local alignment similarity score against the SARS-CoV-2 epitope, using the following parameters: EMBOSS Water, Smith-Waterman algorithm, BLOSUM62 matrix, gap open (10), gap extend (0.5), end gap penalty (false), end gap open penalty (10) and end gap extension penalty (0.5).

### Statistics

Statistical analysis was performed using R v.4.2.3 (96). Data was analyzed with the statistical test described in the corresponding figure legend and annotated with ggsignif v.0.6.4 (97). Pearson or Spearman correlation coefficients and *p*-values were annotated using ggpubr v.0.6.0 (98). Plotting was performed using ggplot2 v.3.4.4 (99). Adjusted *p*-values were obtained by correcting for multiple testing using the Benjamini-Hochberg method, unless otherwise stated. A *p*- or *q*-value < 0.05 was considered statistically significant with non-significant values not displayed. No statistical methods were used to predetermine sample size or power. The experiments were not randomized. Data collection and analysis by investigators were not blinded.

### Data availability

RNA-sequencing gene counts tables, TCR sequences, and predicted epitopes are available as supplemental files. All data, including raw FASTQ files from bulk RNA-sequencing and TCR-sequencing will be made publicly available as part of the larger reference dataset in dbGaP. Some de-identified clinical information from this cohort has been published on PhysioNet (16).

### Code availability

All code used for processing, analysis and figure generation are available at https://github.com/NUPulmonary/2023_Tcell_responses.

## Acknowledgements.

We thank the Robert H. Lurie Comprehensive Cancer Center of Northwestern University in Chicago, IL, for the use of the Flow Cytometry Core Facility. The Lurie Cancer Center is supported in part by an NCI Cancer Center Support Grant #P30 CA060553. This research was supported in part through the computational resources and staff contributions provided for the Quest high performance computing facility at Northwestern University which is jointly supported by the Office of the Provost, the Office for Research, and Northwestern University Information Technology.

The NU SCRIPT Study Investigators are listed in Supplemental File 30.

## Supplemental Table and Figures

**Supplemental Table 1.** Description of the cohort. * Missing values, n: BMI, 1; C-reactive protein, 195; D-dimer, 174; ferritin, 208; lactate, 70; procalcitonin, 114; white blood cell count, 3; absolute neutrophil count, 96; absolute lymphocyte count 101. Empty cells represent data not available or not applicable. BMI, body mass index; SOFA, Sequential Organ Failure Assessment; APS, Acute Physiology Score from APACHE II; LTACH, long-term acute care hospital; SNF, skilled nursing facility; IQR, interquartile range.

|  | Overall | Non-pneumonia control | Other pneumonia | COVID-19 | Other viral pneumonia |
| --- | --- | --- | --- | --- | --- |
| Number | 273 | 33 | 133 | 74 | 33 |
| Age, median [IQR] | 62.0<br>[42.0,71.0] | 62.0<br>[42.0,70.0] | 65.0<br>[51.0,72.0] | 58.5<br>[44.5,66.8] | 60.0<br>[55.0,69.0] |
| Female, n (%) | 108 (39.6) | 16 (48.5) | 55 (41.4) | 23 (31.1) | 14 (42.4) |
| Ethnicity, n (%) |  |  |  |  |  |
| Hispanic or Latino | 60 (22.0) | 5 (15.2) | 13 (9.8) | 37 (50.0) | 5 (15.2) |
| Not Hispanic or Latino | 204 (74.7) | 27 (81.8) | 118 (88.7) | 33 (44.6) | 26 (78.8) |
| Unknown or Not Reported | 9 (3.3) | 1 (3.0) | 2 (1.5) | 4 (5.4) | 2 (6.1) |
| Race, n (%) |  |  |  |  |  |
| Asian | 10 (3.7) | 1 (3.0) | 5 (3.8) | 3 (4.1) | 1 (3.0) |
| Black or African American | 59 (21.6) | 7 (21.2) | 28 (21.1) | 18 (24.3) | 6 (18.2) |
| Unknown or Not Reported | 49 (17.9) | 3 (9.1) | 15 (11.3) | 28 (37.8) | 3 (9.1) |
| White | 155 (56.8) | 22 (66.7) | 85 (63.9) | 25 (33.8) | 23 (69.7) |
| External transfer, n (%) | 91 (33.3) | 12 (36.4) | 43 (32.3) | 24 (32.4) | 12 (36.4) |
| Admission BMI (kg/m2), median [IQR] * | 29.2<br>[24.6,34.3] | 26.4<br>[24.6,32.2] | 27.0<br>[22.1,32.9] | 31.9<br>[28.9,40.6] | 28.3<br>[23.7,31.9] |
| SOFA score on ICU admission, median [IQR] | 11.0 [8.0,13.0] | 11.0 [8.0,13.0] | 11.0 [8.0,14.0] | 11.5 [9.0,13.0] | 10.0 [6.0,14.0] |
| APS score on ICU admission, median [IQR] | 90.0<br>[62.0,108.0] | 80.0<br>[62.0,104.0] | 87.0<br>[64.0,108.0] | 91.0<br>[65.8,109.0] | 83.0<br>[55.0,100.0] |
| Comorbidities, n (%) |  |  |  |  |  |
| Myocardial infarction | 20 (7.3) | 5 (15.2) | 12 (9.0) | 2 (2.7) | 1 (3.0) |
| Congestive heart failure | 81 (29.7) | 13 (39.4) | 50 (37.6) | 12 (16.2) | 6 (18.2) |
| Peripheral vascular disease | 63 (23.1) | 8 (24.2) | 39 (29.3) | 9 (12.2) | 7 (21.2) |
| Cerebrovascular disease | 57 (20.9) | 6 (18.2) | 32 (24.1) | 8 (10.8) | 11 (33.3) |
| Dementia | 13 (4.8) | 3 (9.1) | 4 (3.0) | 2 (2.7) | 4 (12.1) |
| Chronic pulmonary disease | 96 (35.2) | 13 (39.4) | 53 (39.8) | 17 (23.0) | 13 (39.4) |
| Rheumatic disease | 21 (7.7) | 4 (12.1) | 11 (8.3) | 3 (4.1) | 3 (9.1) |
| Peptic ulcer disease | 27 (9.9) | 5 (15.2) | 16 (12.0) | 1 (1.4) | 5 (15.2) |
| Liver disease | 80 (29.3) | 12 (36.4) | 44 (33.1) | 11 (14.9) | 13 (39.4) |
| Diabetes | 104 (38.1) | 10 (30.3) | 41 (30.8) | 35 (47.3) | 18 (54.5) |
| Hemiplegia or paraplegia | 19 (7.0) | 3 (9.1) | 12 (9.0) | 2 (2.7) | 2 (6.1) |
| Renal disease | 77 (28.2) | 14 (42.4) | 41 (30.8) | 11 (14.9) | 11 (33.3) |
| Cancer | 87 (31.9) | 12 (36.4) | 48 (36.1) | 11 (14.9) | 16 (48.5) |
| Immunocompromise | 70 (25.6) | 8 (24.2) | 40 (30.1) | 6 (8.1) | 16 (48.5) |
| Biomarkers on day of first BAL procedure, median [IQR] |  |  |  |  |  |
| C-reactive protein, mg/dL * | 163.2<br>[92.0,259.5] | 12.0<br>[11.0,55.0] | 96.0<br>[22.5,176.0] | 169.0<br>[116.0,275.5] | 168.0<br>[168.0,168.0] |
| D-dimer, ng/mL * | 1103.0<br>[492.5,3540.5] | 1345.5<br>[405.8,2220.5] | 2922.0<br>[886.0,5091.0] | 896.5<br>[468.8,2892.6] | 1954.2<br>[1170.5,2931.1] |
| Ferritin, ng/mL * | 663.2<br>[300.9,1210.3] | 92.4<br>[57.0,191.2] | 631.8<br>[319.8,1183.7] | 766.4<br>[330.1,1355.2] | — |
| Lactate, mmol/L * | 1.5 [1.1,2.1] | 1.7 [1.1,2.3] | 1.6 [1.1,2.4] | 1.4 [1.0,1.8] | 1.3 [1.0,1.7] |
| Procalcitonin, ng/mL * | 0.6 [0.2,2.8] | 0.4 [0.1,1.6] | 0.6 [0.3,2.9] | 0.4 [0.1,2.9] | 0.8 [0.3,3.8] |
| White blood cell count, per 1,000/μL * | 11.6 [8.0,16.7] | 12.6 [8.1,20.4] | 12.6 [8.7,16.8] | 10.1 [7.1,14.6] | 10.7 [7.2,18.5] |
| Absolute neutrophil count, per 1,000/μL * | 9.2 [5.7,13.7] | 12.2 [6.0,18.8] | 9.6 [6.3,15.3] | 8.7 [5.5,11.2] | 8.9 [5.2,13.0] |
| Absolute lymphocyte count, per 1,000/μL * | 0.8 [0.5,1.4] | 0.6 [0.4,0.8] | 0.9 [0.5,1.7] | 1.0 [0.6,1.3] | 0.8 [0.2,1.6] |
| Medications |  |  |  |  |  |
| Corticosteroids administered during admission, n (%) | 162 (59.3) | 20 (60.6) | 80 (60.2) | 39 (52.7) | 23 (69.7) |
| Prednisone equivalents administered during admission, mg | 70.0<br>[0.0,240.0] | 80.0<br>[0.0,340.0] | 92.0<br>[0.0,250.0] | 15.0<br>[0.0,209.0] | 96.0<br>[0.0,272.0] |
| Tocilizumab, n (%) | 13 (4.8) | — | 1 (0.8) | 12 (16.2) | — |
| Sarilumab, n (%) | 12 (4.4) | — | — | 12 (16.2) | — |
| Remdesivir, n (%) | 12 (4.4) | — | — | 12 (16.2) | — |
| Remdesivir or placebo, n (%) | 9 (3.3) | — | — | 9 (12.2) | — |
| ICU course |  |  |  |  |  |
| Ventilation duration (days), median [IQR] | 10.0<br>[4.0,22.0] | 3.0<br>[2.0,9.0] | 9.0<br>[4.0,19.0] | 20.0<br>[10.0,33.0] | 7.0<br>[3.0,13.0] |
| ICU length of stay (days), median [IQR] | 13.0<br>[6.0,23.0] | 6.0<br>[4.0,11.0] | 11.0<br>[5.0,21.0] | 21.0<br>[14.0,35.0] | 11.0<br>[9.0,20.0] |
| Tracheostomy, n (%) | 70 (25.6) | 3 (9.1) | 29 (21.8) | 32 (43.2) | 6 (18.2) |
| Number of BALs, n (% of total) | 432 (100) | 36 (8.3) | 187 (43.3) | 165 (38.2) | 44 (10.2) |
| Discharge disposition, n (%) |  |  |  |  |  |
| Died | 86 (31.5) | 10 (30.3) | 47 (35.3) | 19 (25.7) | 10 (30.3) |
| Home | 76 (27.8) | 12 (36.4) | 27 (20.3) | 30 (40.5) | 7 (21.2) |
| LTACH | 35 (12.8) | 3 (9.1) | 17 (12.8) | 12 (16.2) | 3 (9.1) |
| Rehab | 50 (18.3) | 4 (12.1) | 27 (20.3) | 9 (12.2) | 10 (30.3) |
| SNF | 18 (6.6) | 2 (6.1) | 12 (9.0) | 4 (5.4) | — |
| Hospice | 8 (2.9) | 2 (6.1) | 3 (2.3) | 0 (0.0) | 3 (9.1) |

**Supplemental Figure 1.**
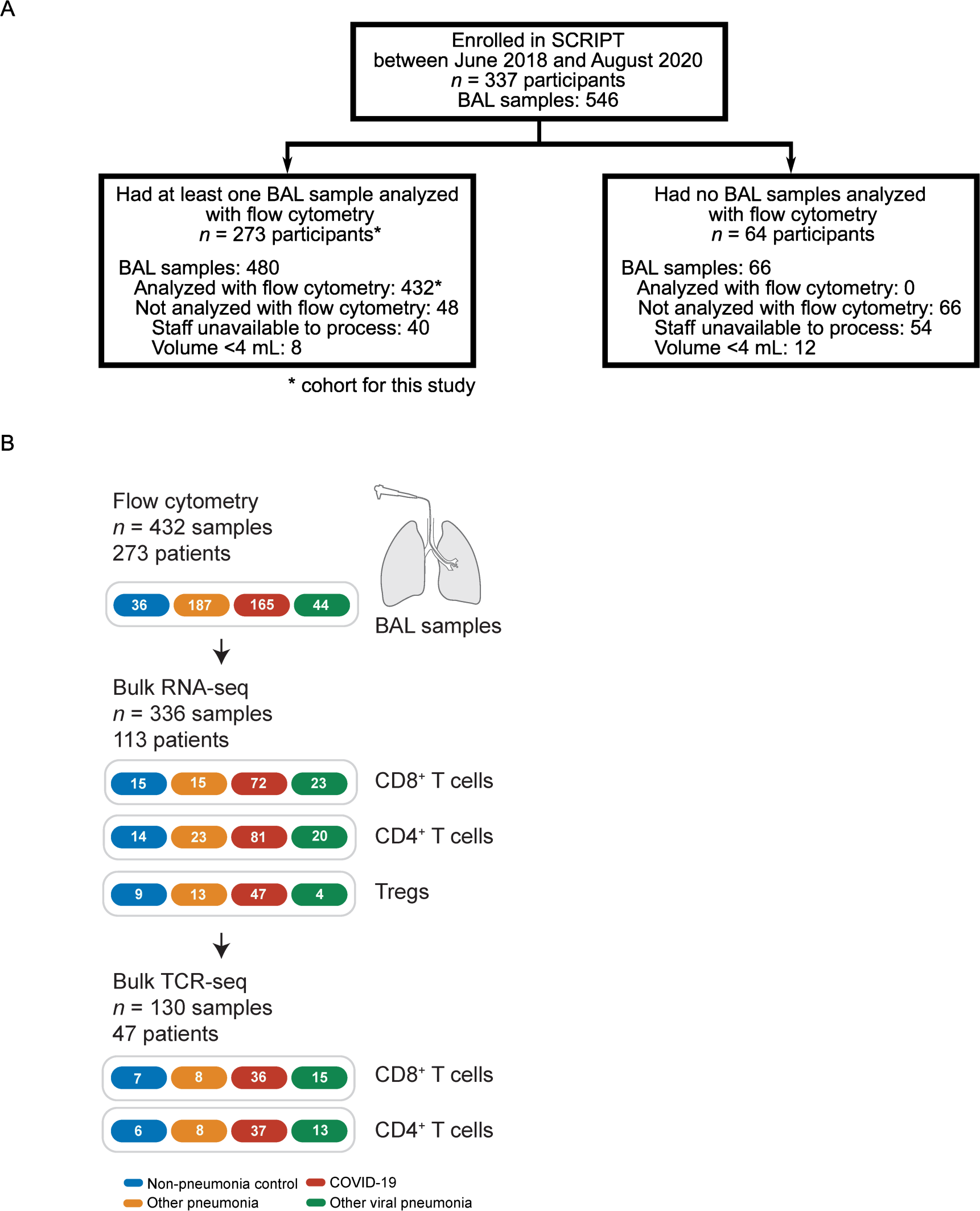
**(A)** CONSORT diagram of patients included in this study. **(B)** Schematic depicting multi-step analysis of BAL fluid samples with flow cytometry, bulk RNA-sequencing, and bulk TCR-sequencing by diagnosis and T cell subset.

**Supplemental Figure 2.**
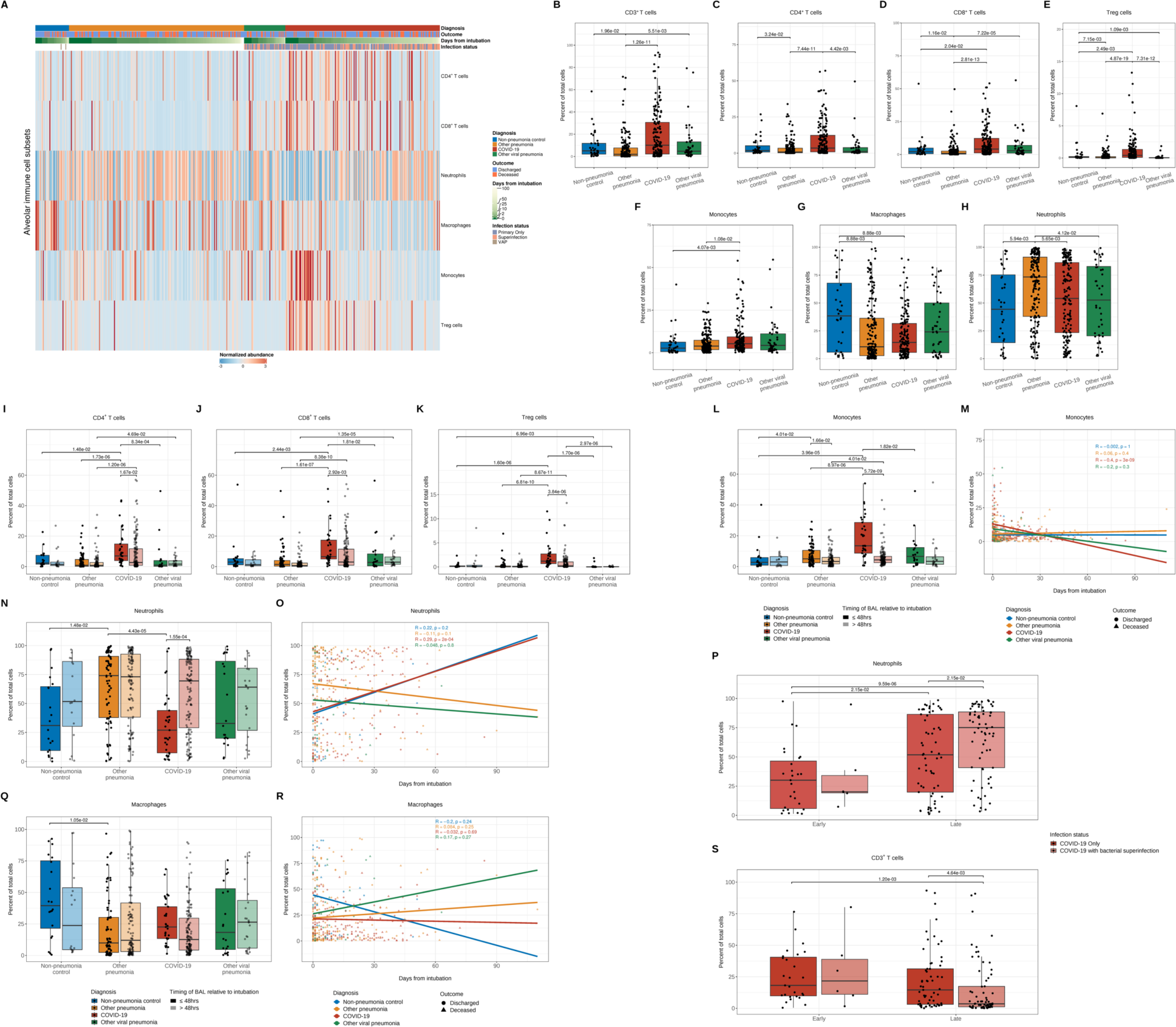
SARS-CoV-2 pneumonia is characterized by a lymphomonocytic alveolar infiltrate early following intubation. **(A)** Heatmap of flow cytometry analysis of alveolar immune cell subsets from BAL fluid samples ordered by duration of mechanical ventilation (blanks indicate chronically ventilated patients) and grouped by diagnosis, binary outcome (whether a given patient was discharged or died during hospitalization), and infection status (presence or absence of bacterial superinfection in patients with COVID-19 or other viral pneumonia). Blanks in these two groups refer to samples for which microbiological data were incomplete and infectious status could not be determined. The VAP (ventilator-associated pneumonia) flag designates samples from non-pneumonia controls or patients with COVID-19 or other viral pneumonia who cleared the virus and then developed a bacterial pneumonia. **(B-H)** Percent of alveolar immune cell subsets detected in BAL fluid samples from flow cytometry analysis (*q* < 0.05, pairwise Wilcoxon rank-sum tests with FDR correction). **(I-L**, **N** and **Q)** Comparison of alveolar immune cell subset percentages between early (≤48 hours following intubation) and late (>48 hours following intubation) samples (*q* < 0.05, pairwise Wilcoxon rank-sum tests with FDR correction). **(M, O** and **R)** Correlation analysis between the percentage of alveolar immune cell subsets and duration of mechanical ventilation with Pearson correlation coefficient. **(P** and **S)** Comparison of neutrophil (P) and CD3^+^ T cell (S) percentage grouped by the presence or absence of bacterial superinfection in early (≤48 hours following intubation) and late (>48 hours following intubation) COVID-19 samples (*q* < 0.05, pairwise Wilcoxon rank-sum tests with FDR correction).

**Supplemental Figure 3.**
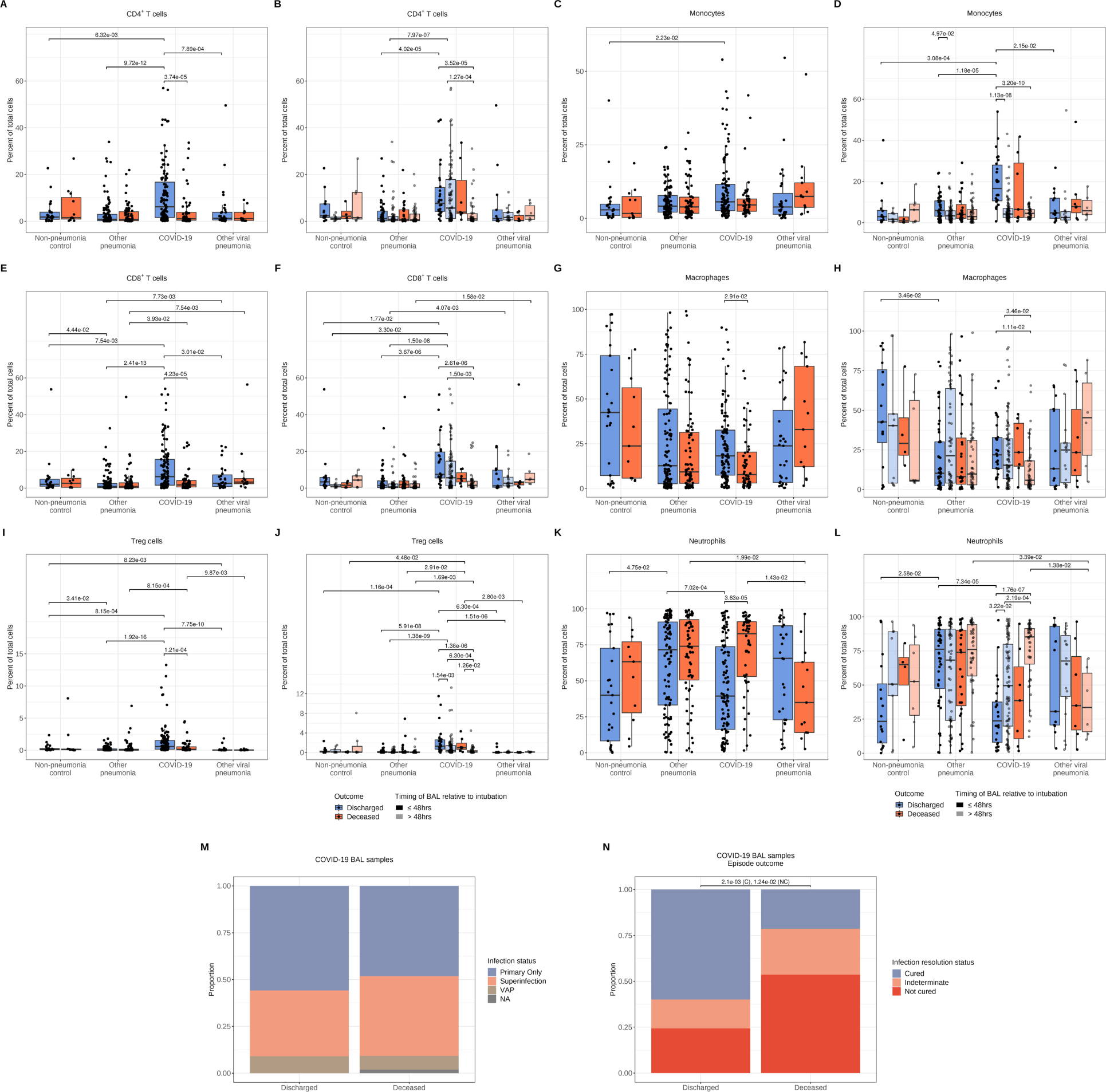
Persistent alveolar T cell enrichment throughout the course of severe SARS-CoV-2 pneumonia is associated with discharge from hospital. **(A-L)** Comparison of alveolar immune cell subset percentages by binary outcome (C, E, G, I, and K) and between timing of BAL sampling (B, D, F, H, J and L) (*q* <0.05, pairwise Wilcoxon rank-sum tests with FDR correction). **(M)** Proportion of BAL fluid samples from patients with COVID-19, comparing presence or absence of bacterial superinfection with binary outcome (not significant by Fisher exact test). **(N)** Proportion of BAL fluid samples from patients with COVID-19, comparing pneumonia episode outcome status with binary outcome (*q* < 0.05, Fisher exact test with FDR correction). C (Cured) and NC (Not cured).

**Supplemental Figure 4.**
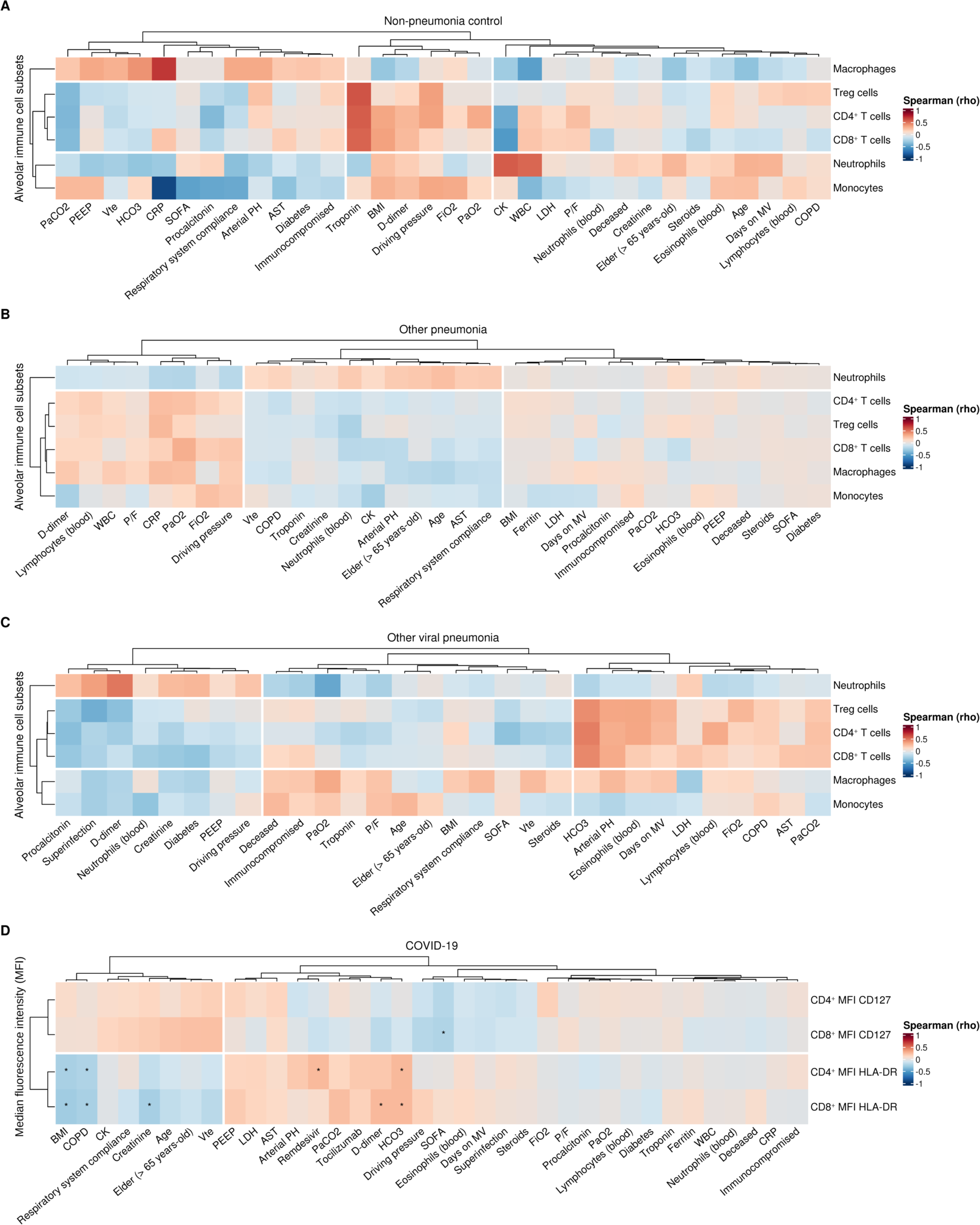
Correlation of alveolar immune cell subset abundance with clinical outcomes differs between patients with distinct etiologies of severe pneumonia. **(A-C).** Correlation analysis between the percentage of alveolar immune cell subsets and clinical, physiologic, and laboratory variables as a function of diagnostic group. No significant values after calculating Spearman rank correlation coefficient with FDR correction. **(D)** Correlation between T cell subset surface expression of CD127 and HLA-DR in the alveolar space with clinical, laboratory, and physiological variables in COVID-19 samples. Spearman rank correlation coefficient with FDR correction (*q* < 0.05 [*]). Abbreviations: PaCO2 (partial arterial carbon dioxide pressure), HCO3 (bicarbonate), Days on MV (days on mechanical ventilation), SOFA (Sequential Organ Failure Assessment), WBC (peripheral white blood cells), CK (creatinine kinase), Vte (minute ventilation), LDH (lactate dehydrogenase), FiO2 (fraction of inspired oxygen), CRP (C-reactive protein), PEEP (positive end-expiratory pressure), BMI (body mass index), AST (aspartate aminotransferase), PaO2 (partial arterial oxygen pressure), P/F (ratio of partial arterial oxygen pressure to fraction of inspired oxygen), COPD (chronic obstructive pulmonary disease), MFI (median fluorescence intensity).

**Supplemental Figure 5.**
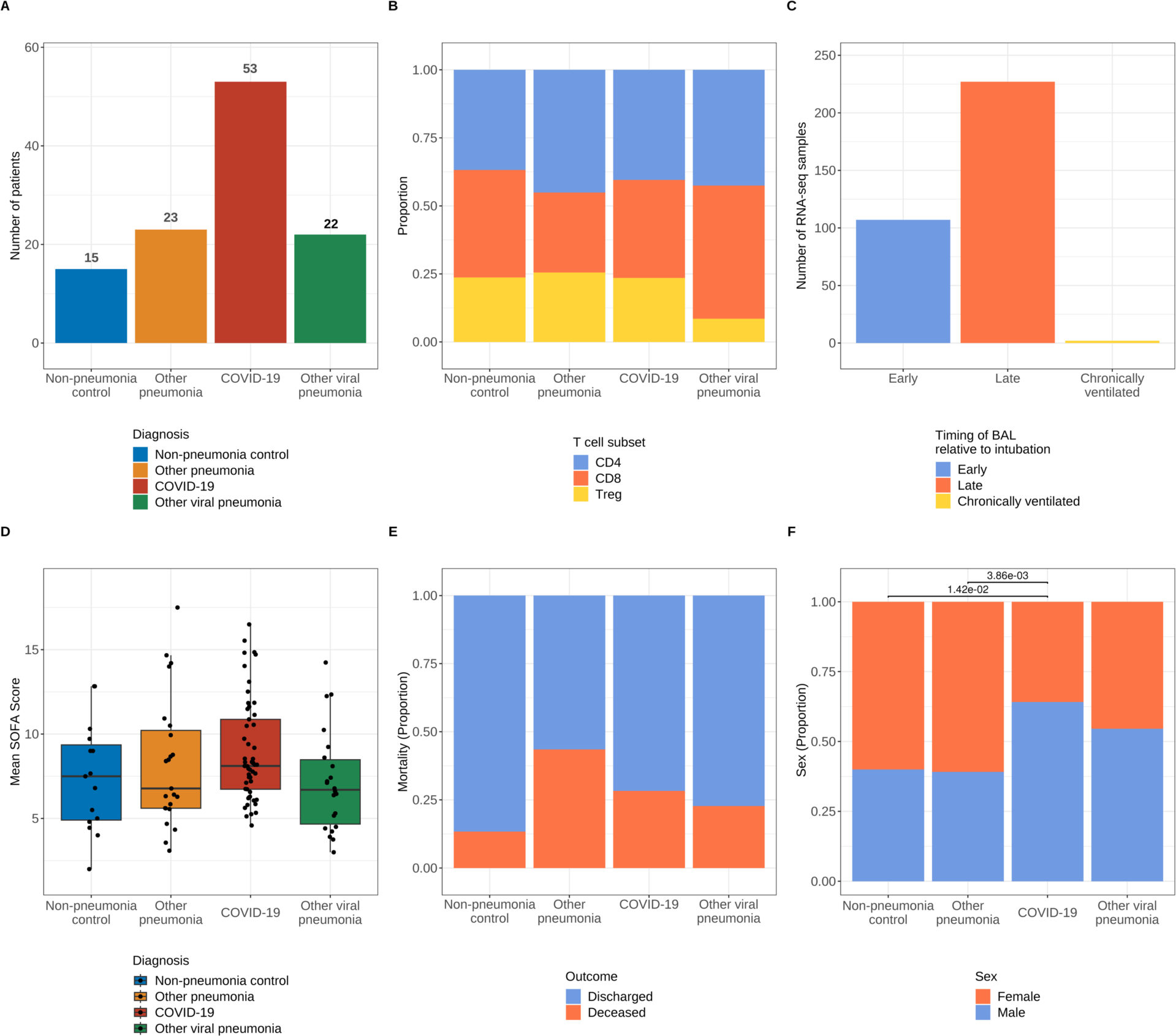
Composition and demographics of bulk RNA-sequencing samples. **(A)** Number of participants grouped by diagnosis. **(B)** Proportion of samples grouped by T cell subset and diagnosis. **(C)** Number of samples categorized by timing of BAL. **(D)** Distribution of Sequential Organ Failure Assessment (SOFA) scores by patient. Nonsignificant after pairwise Wilcoxon rank-sum tests with FDR correction). **(E)** Mortality by patient. Nonsignificant after pairwiseχ￼ ^2^ tests for homogeneity of proportions with FDR correction. **(F)** Sex by patient (pairwiseχ￼ ^2^ tests for homogeneity of proportions with FDR correction).

**Supplemental Figure 6.**
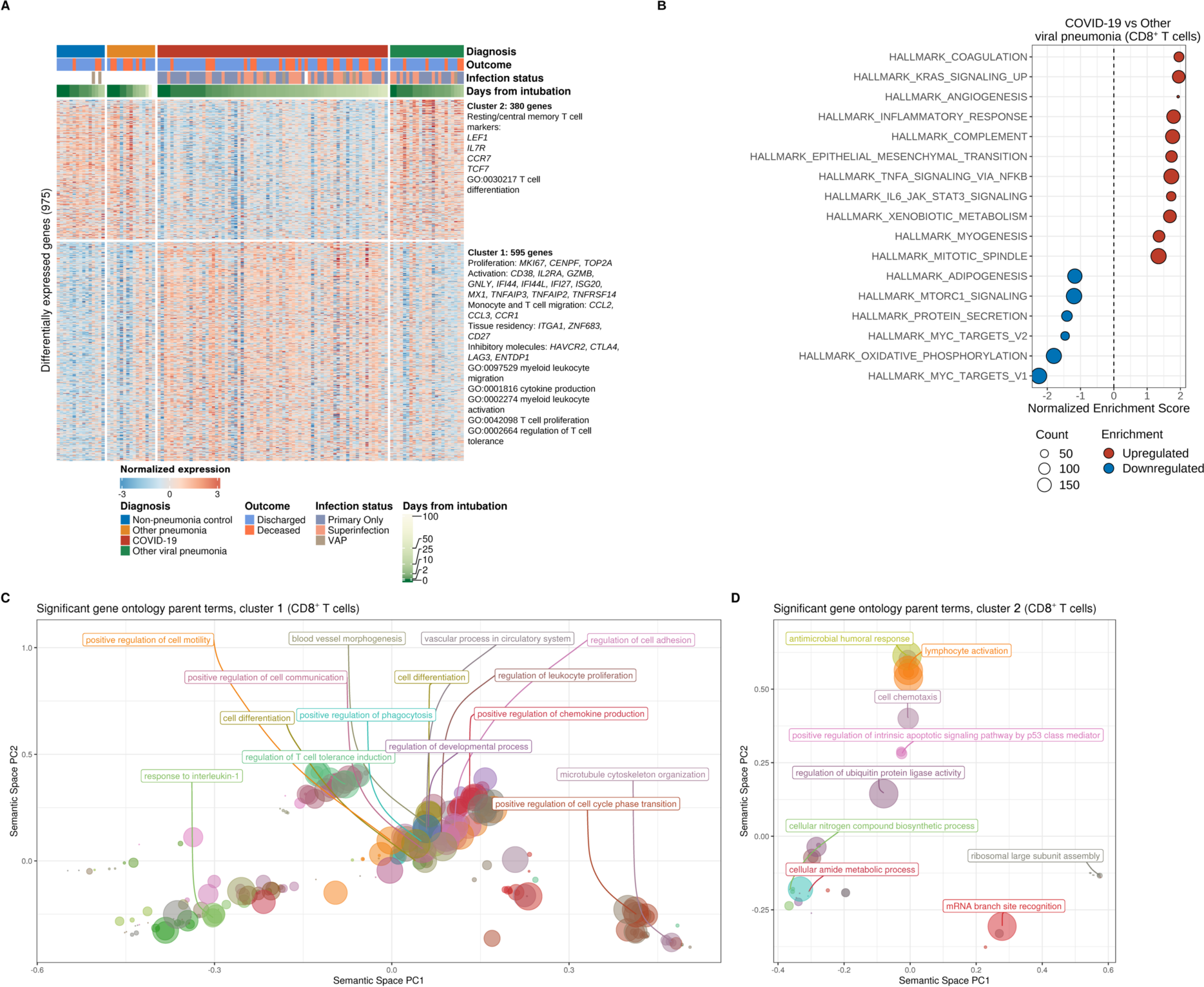
SARS-CoV-2 pneumonia is characterized by a transcriptional program enriched for processes associated with monocyte and T cell activation, migration, and angiogenesis in alveolar CD8^+^ T cells. (A) *K*-means clustering of 975 differentially expressed genes (*q* < 0.05, likelihood-ratio test with FDR correction) across pneumonia diagnoses. Columns represent unique samples grouped by diagnosis and are ordered by duration of mechanical ventilation. Column headers are color-coded by diagnosis, binary outcome (whether a given patient was discharged or died during hospitalization), duration of mechanical ventilation (blanks indicate chronically ventilated patients), and infection status (presence or absence of bacterial superinfection in patients with COVID-19 or other viral pneumonia). The VAP (ventilator-associated pneumonia) flag designates samples from non-pneumonia controls or patients with COVID-19 or other viral pneumonia who cleared the virus and then developed a bacterial pneumonia. Representative genes and significant gene ontology (GO) biological processes are shown for each cluster. **(B)** Gene set enrichment analysis (GSEA) of Hallmark gene sets for the pairwise comparison between COVID-19 samples and other viral pneumonia samples. Count denotes pathway size after removing genes not detected in the expression dataset. Enrichment denotes significant (*q* < 0.25 with FDR correction) upregulated (red) and downregulated (blue) pathways by normalized enrichment score. **(C-D)** Gene ontology (GO) parent term annotation after grouping significant terms (following classical or over-representation enrichment analysis, *q* < 0.05 with multiple testing correction using the Benjamini-Hochberg method) by semantic similarity. Reduced GO terms are depicted in a scatter plot where distance between points represent the similarity between terms and axes are the first two components after applying a principal coordinates analysis to the dissimilarity matrix. Points are color-coded by unique terms and size denotes the number of genes within each GO term.

**Supplemental Figure 7.**
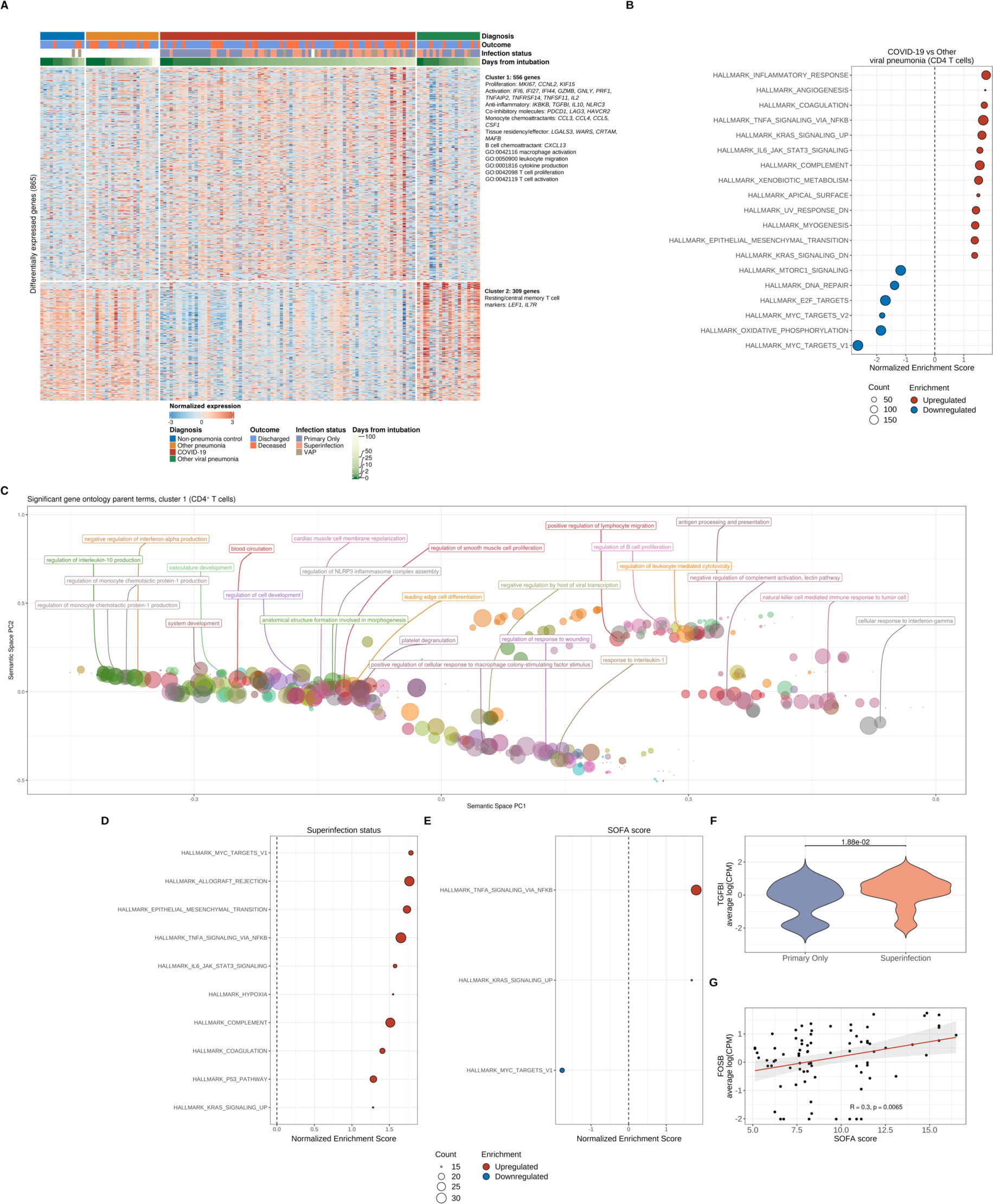
SARS-CoV-2 pneumonia is characterized by a transcriptional program enriched for processes associated with monocyte, B cell and T cell activation, migration, and angiogenesis in alveolar CD4^+^ T cells. **(A)** *K*-means clustering of 865 differentially expressed genes (*q* < 0.05, likelihood-ratio test with FDR correction) across pneumonia diagnoses. Columns represent unique samples grouped by diagnosis and are ordered by duration of mechanical ventilation. Column headers are color-coded by diagnosis, binary outcome (whether a given patient was discharged or died during hospitalization), duration of mechanical ventilation (blanks indicate chronically ventilated patients), and infection status (presence or absence of bacterial superinfection in patients with COVID-19 or other viral pneumonia). The VAP (ventilator-associated pneumonia) flag designates samples from non-pneumonia controls or patients with COVID-19 or other viral pneumonia who cleared the virus and then developed a bacterial pneumonia. Representative genes and significant gene ontology (GO) biological processes are shown for each cluster. **(B)** Gene set enrichment analysis (GSEA) of Hallmark gene sets for the pairwise comparison between COVID-19 samples and other viral pneumonia samples. Count denotes pathway size after removing genes not detected in the expression dataset. Enrichment denotes significant (*q* < 0.25 with FDR correction) upregulated (red) and downregulated (blue) pathways by normalized enrichment score. **(C)** Gene ontology (GO) parent term annotation after grouping significant terms (following classical or over-representation enrichment analysis, *q* < 0.05 with multiple testing correction using the Benjamini-Hochberg method) by semantic similarity. Reduced GO terms are depicted in a scatter plot where distance between points represent the similarity between terms and axes are the first two components after applying a principal coordinates analysis to the dissimilarity matrix. Points are color-coded by unique terms and size denotes the number of genes within each GO term. **(D-E)** GSEA of COVID-19 samples after performing correlation analysis of differentially expressed genes in CD4^+^ T cells and clinical variables of interest with Spearman rank correlation coefficient computation. Count denotes pathway size after removing genes not detected in the expression dataset. Enrichment denotes significant (*q* < 0.25 with FDR correction) upregulated (red) and downregulated (blue) pathways by normalized enrichment score. **(F-G)** Leading edge analysis reveals selected core genes driving pathway enrichment signal in clinical variables, which are annotated for the superinfection and SOFA score variables.

**Supplemental Figure 8.**
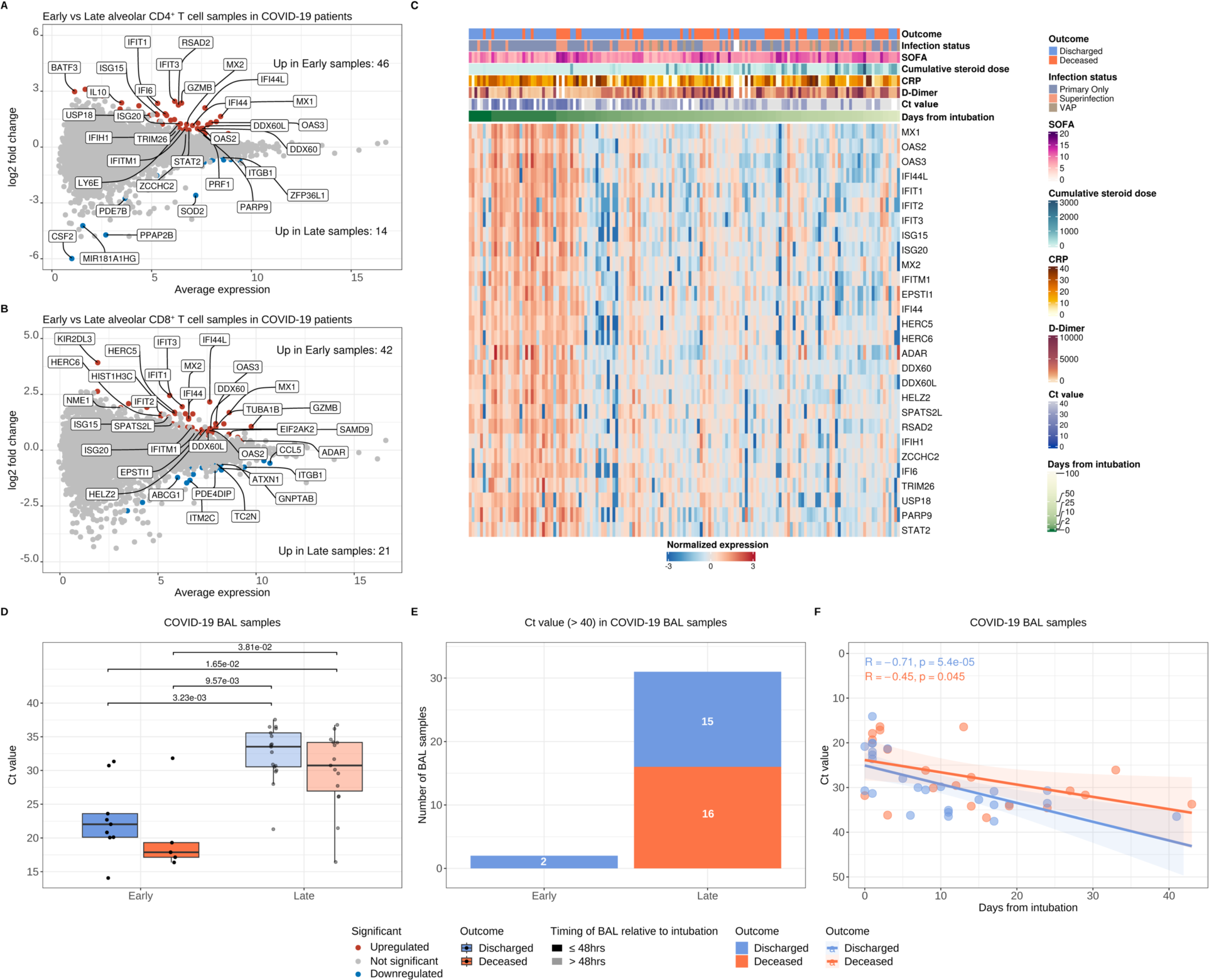
The early T cell response during severe SARS-CoV-2 pneumonia is dominated by an interferon signaling transcriptional program. **(A-B)** MA plot of differentially expressed genes in CD4^+^ T cells (81 samples from 46 patients with COVID-19 pneumonia) (A) and CD8^+^ T cells (72 samples from 46 patients with COVID-19 pneumonia) (B), comparing early (≤48 hours following intubation) versus late (>48 hours following intubation) COVID-19 samples. Significantly upregulated genes in early samples are shown in red, and significantly upregulated genes in late samples are shown in blue (*q* < 0.05, likelihood-ratio tests with FDR correction). Genes shown in gray are not significantly differentially expressed. Representative significant genes are annotated. **(C)** Heatmap of longitudinal analysis of interferon-stimulated genes in combined CD4^+^ and CD8^+^ T cells of patients with severe SARS-CoV-2 pneumonia. Columns represent unique T cell samples and are color-coded by binary outcome, infection status, severity of illness (SOFA score), cumulative steroid dose (mg of hydrocortisone), C reactive protein (CRP), D-dimer, viral load (Ct value), and ordered by duration of mechanical ventilation. Blanks indicate missing values. **(D)** Comparison of SARS-CoV-2 viral load (Ct value) by binary outcome and BAL sampling time in COVID-19 samples that underwent RNA-sequencing (*q* < 0.05, pairwise Wilcoxon rank-sum tests with FDR correction). **(E)** Number of COVID-19 BAL samples grouped by binary outcome and sampling time with a Ct value above limit of detection (>40). **(F)** Correlation analysis of COVID-19 Ct values grouped by binary outcome and duration of mechanical ventilation with Spearman rank correlation coefficient.

**Supplemental Figure 9.**
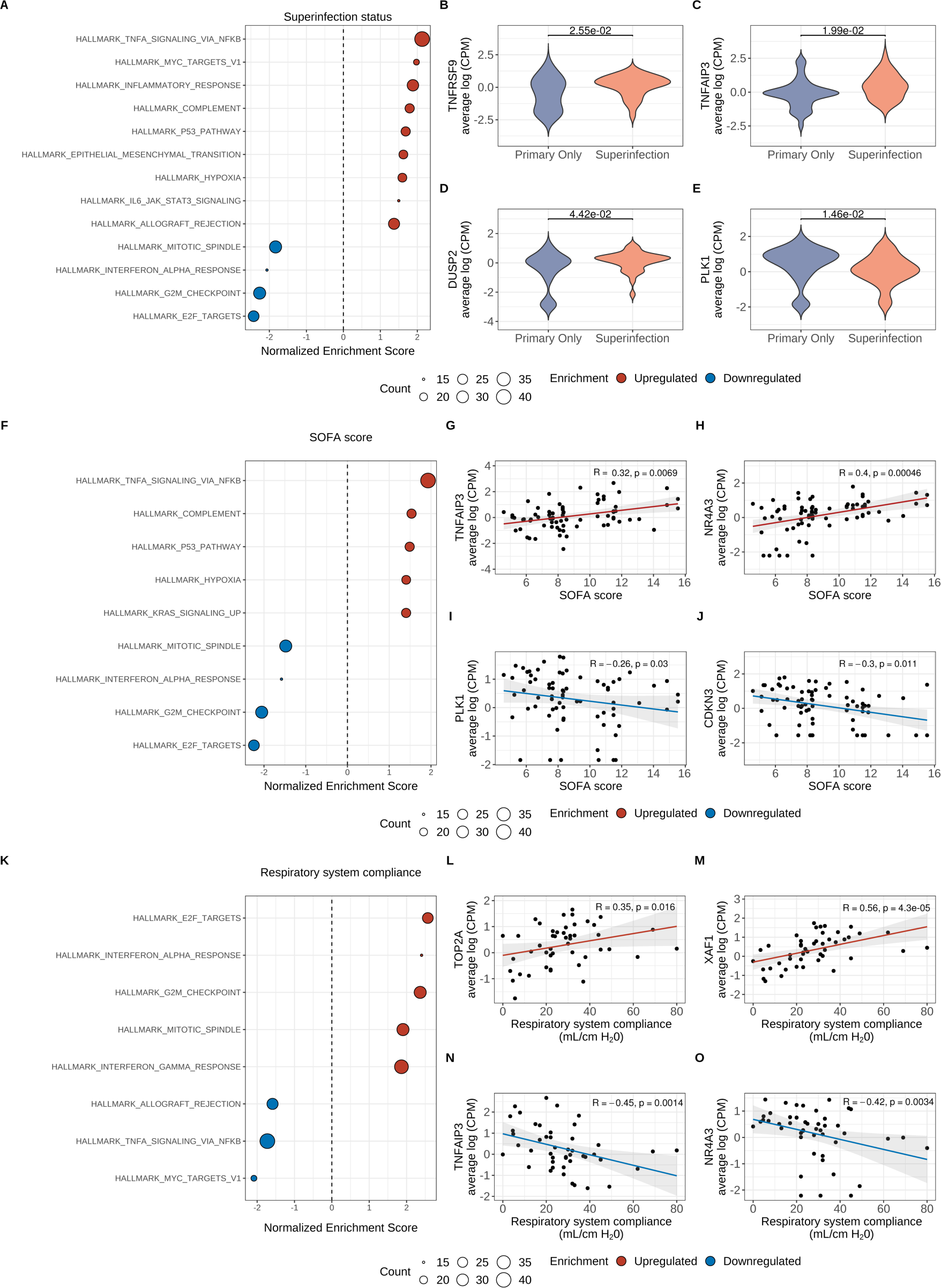
Distinct activation molecular signatures in alveolar CD8^+^ T cells predict clinical outcomes in patients with severe SARS-CoV-2 pneumonia. **(A-O)** Gene set enrichment analysis (GSEA) of COVID-19 samples after performing correlation analysis of differentially expressed genes in CD8^+^ T cells and clinical variables of interest with Spearman rank correlation coefficient computation. Count denotes pathway size after removing genes not detected in the expression dataset. Enrichment denotes significant (*q* < 0.25 with FDR correction) upregulated (red) and downregulated (blue) pathways by normalized enrichment score. Leading edge analysis reveals selected core genes driving pathway enrichment signal in clinical variables, which are annotated for the superinfection (B-E), severity of illness (SOFA score) (G-J), and respiratory system compliance (L-O) variables.

**Supplemental Figure 10.**
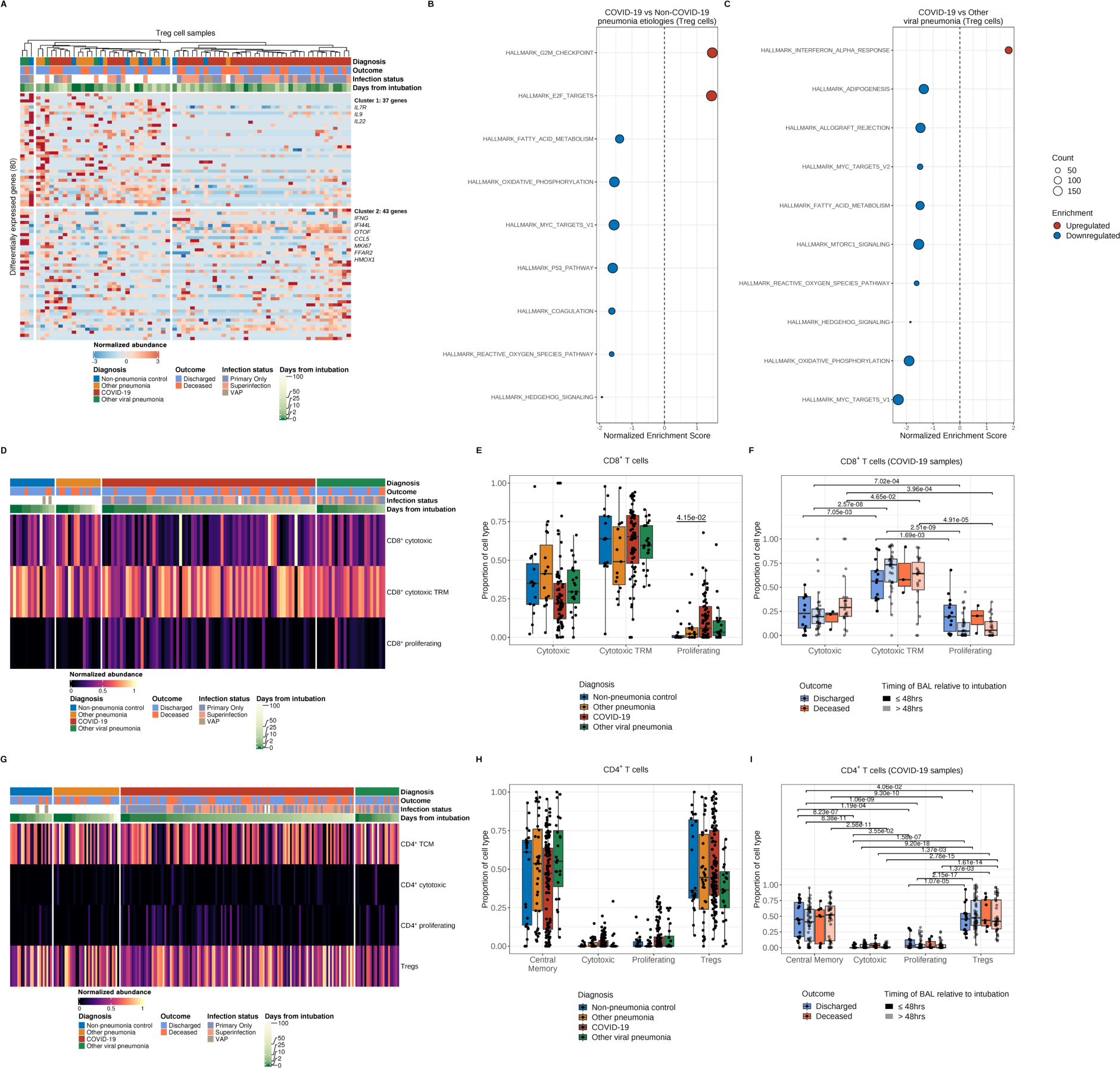
Cell-type deconvolution of bulk RNA-sequencing reveals a dominant activated memory phenotype of alveolar T cells during severe pneumonia. **(A)** *K*-means clustering of 80 differentially expressed genes (*q* < 0.05, likelihood-ratio test with FDR correction) in Treg cells across pneumonia diagnoses. Columns represent unique samples and column headers are color-coded by diagnosis, binary outcome (whether a given patient was discharged or died during hospitalization), duration of mechanical ventilation (blanks indicate chronically ventilated patients), and infection status (presence or absence of bacterial superinfection in patients with COVID-19 or other viral pneumonia). The VAP (ventilator-associated pneumonia) flag designates samples from non-pneumonia controls or patients with COVID-19 or other viral pneumonia who cleared the virus and then developed a bacterial pneumonia. Samples were clustered using Ward’s minimum variance clustering method. Representative genes are shown for each cluster. **(B-C)** Gene set enrichment analysis (GSEA) of Hallmark gene sets for the pairwise comparison between COVID-19 samples and combined non-COVID-19 samples (non-pneumonia control, other pneumonia, and other viral pneumonia) (B) or the pairwise comparison between COVID-19 samples and other viral pneumonia samples (C). Count denotes pathway size after removing genes not detected in the expression dataset. Enrichment denotes significant (*q* < 0.25 with FDR correction) upregulated (red) and downregulated (blue) pathways by normalized enrichment score. **(D)** Heatmap demonstrating the proportion of alveolar CD8^+^ T cell subsets from deconvolution analysis. Columns represent unique samples grouped by diagnosis and are ordered by duration of mechanical ventilation. Column headers are color-coded by diagnosis, binary outcome, duration of mechanical ventilation (blanks indicate chronically ventilated patients), and infection status (presence or absence of bacterial superinfection in patients with COVID-19 or other viral pneumonia). The VAP (ventilator-associated pneumonia) flag designates samples from non-pneumonia controls or patients with COVID-19 or other viral pneumonia who cleared the virus and then developed a bacterial pneumonia. **(E)** Proportion of alveolar CD8^+^ T cell subsets across different pneumonia etiologies. (*q* < 0.05, pairwise Wilcoxon rank-sum tests with FDR correction). **F)** Proportion of alveolar CD8^+^ T cell subsets by outcome and timing of BAL fluid sampling in COVID-19 patients (*q* < 0.05, pairwise Wilcoxon rank-sum tests with FDR correction). **(G)** Heatmap demonstrating the proportion of alveolar CD4^+^ T cell subsets from deconvolution analysis. Columns represent unique samples grouped by diagnosis and are ordered by duration of mechanical ventilation. Column headers are color-coded by diagnosis, binary outcome, duration of mechanical ventilation (blanks indicate chronically ventilated patients), and infection status (presence or absence of bacterial superinfection in patients with COVID-19 or other viral pneumonia). The VAP (ventilator-associated pneumonia) flag designates samples from non-pneumonia controls or patients with COVID-19 or other viral pneumonia who cleared the virus and then developed a bacterial pneumonia. **(H)** Proportion of alveolar CD4^+^ T and Treg cell subsets across different pneumonia etiologies (*q* < 0.05, pairwise Wilcoxon rank-sum tests with FDR correction). **(I)** Proportion of alveolar CD4^+^ T cell subsets by binary outcome and timing of BAL fluid sampling in COVID-19 patients (*q* < 0.05, pairwise Wilcoxon rank-sum tests with FDR correction).

**Supplemental Figure 11.**
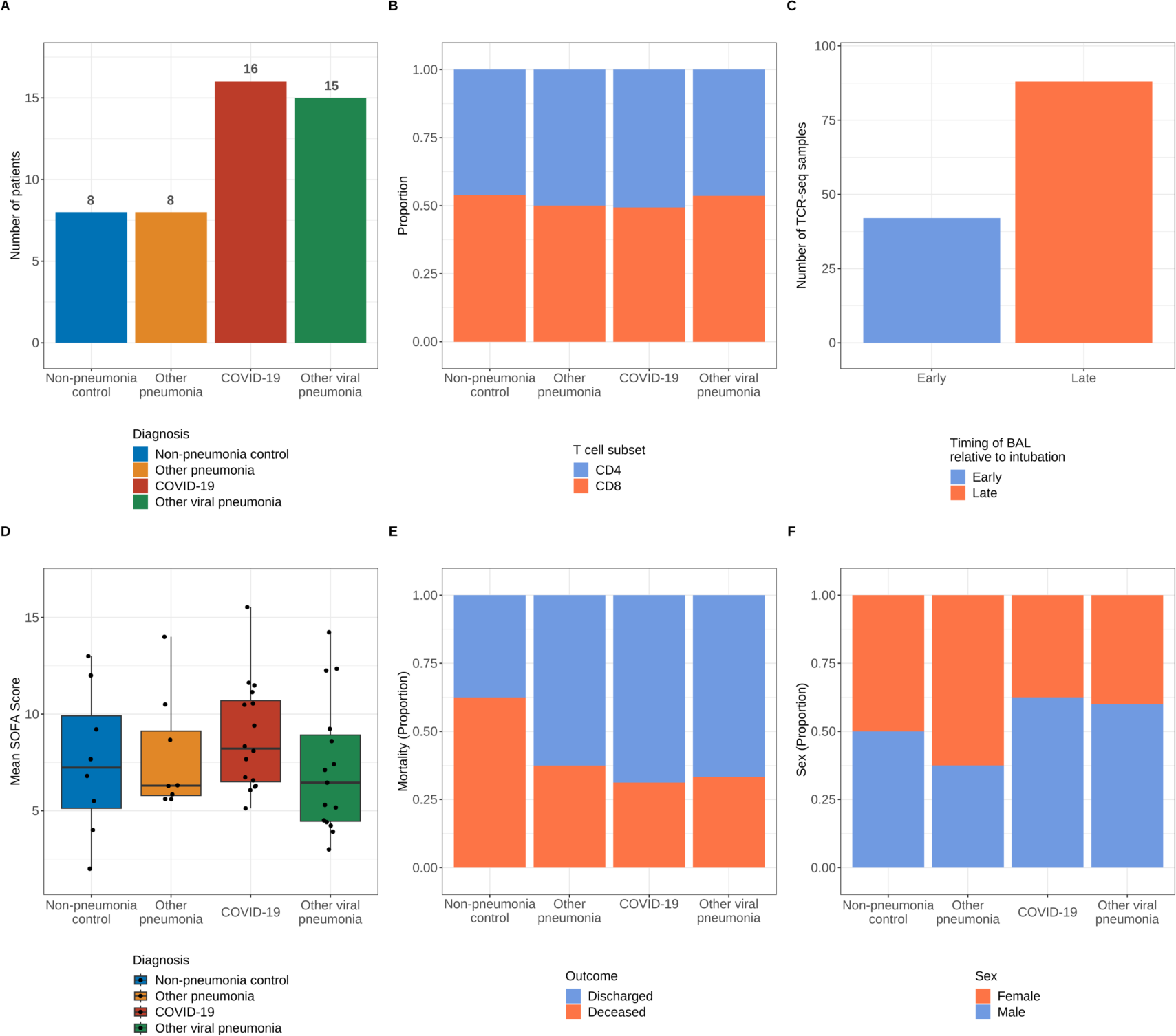
Composition and demographics of bulk TCR-sequencing samples. **(A)** Number of participants grouped by diagnosis. **(B)** Proportion of samples grouped by T cell subset and diagnosis. **(C)** Number of samples categorized by timing of BAL. **(D)** Distribution of SOFA scores by patient. Nonsignificant after pairwise Wilcoxon rank-sum tests with FDR correction. **(E)** Mortality by patient. Nonsignificant after pairwise χ^2^ tests for homogeneity of proportions with FDR correction. **(F)** Sex by patient. Nonsignificant after pairwise χ^2^ tests for homogeneity of proportions with FDR correction.

**Supplemental Figure 12.**
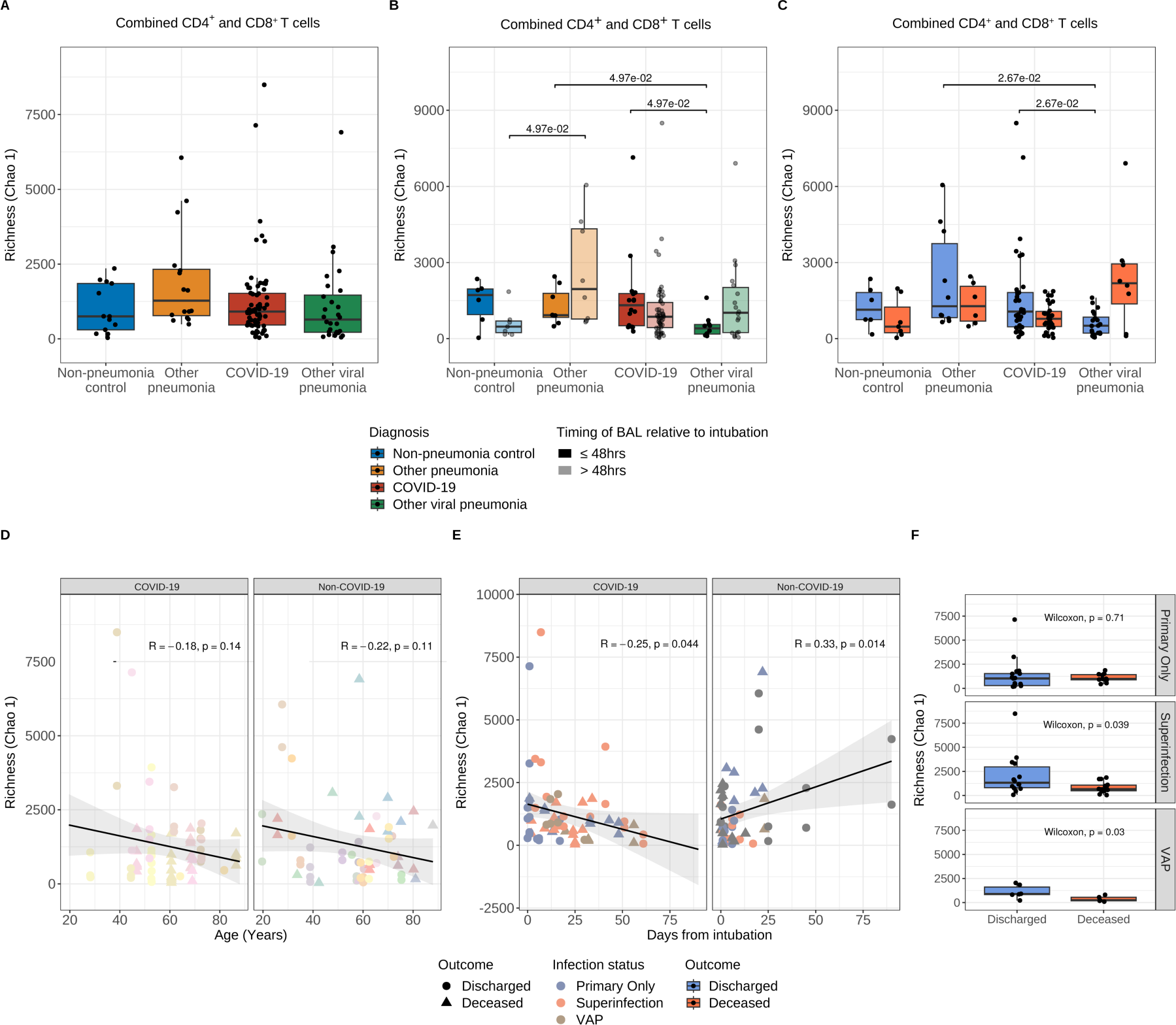
SARS-CoV-2 pneumonia complicated by secondary bacterial pneumonia is associated with lower TCR repertoire diversity. **(A-C)** Alpha diversity estimation with Chao 1 in combined alveolar CD4^+^ and CD8^+^ T cells in patients grouped by diagnosis (A), timing of BAL fluid collection relative to intubation (B), and binary outcome (C) (*q* < 0.05, pairwise Wilcoxon rank-sum tests with FDR correction). **(D-E)** Correlation analysis between combined alveolar CD4^+^ and CD8^+^ T cell richness (Chao 1) and age (D) and duration of mechanical ventilation (E) using Pearson correlation. Data points are color-coded by unique patients (D) or infection status (E) and shaped according to binary outcome. **(F)** Alpha diversity estimation with Chao1 in combined alveolar CD8^+^ and CD4^+^ T cells in patients grouped by binary outcome and infection status (presence or absence of bacterial superinfection in patients with COVID-19 or other viral pneumonia). The VAP (ventilator-associated pneumonia) flag designates samples from non-pneumonia controls or patients with COVID-19 or other viral pneumonia who cleared the virus and then developed a bacterial pneumonia. Wilcoxon rank sum tests *p*-values are shown.

**Supplemental Figure 13.**
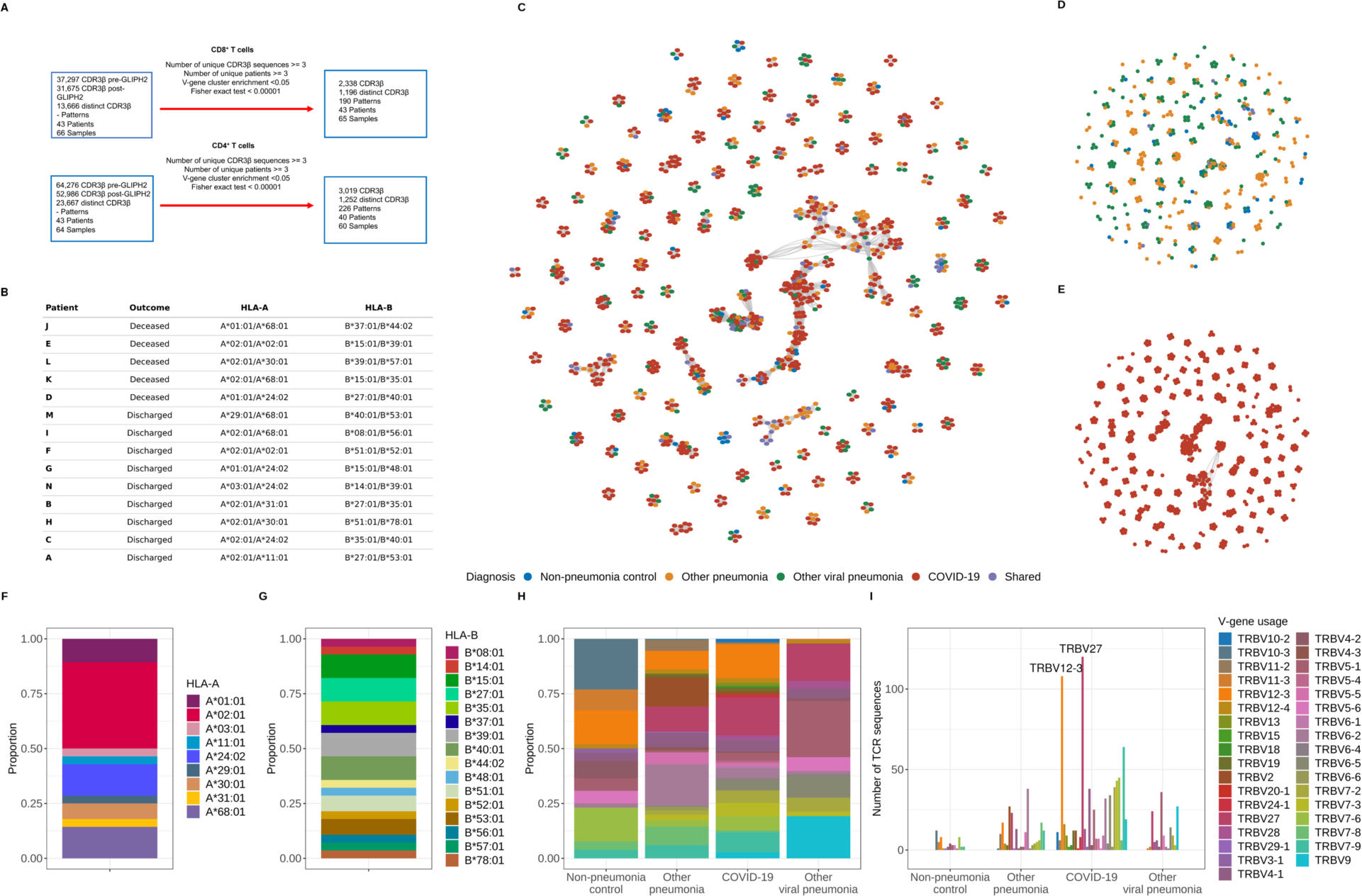
Alveolar CD8^+^ T cell receptor specificity analysis in SARS-CoV-2 pneumonia. **(A)** GLIPH2 specificity filtering criteria and numbers. **(B)** Inferred HLA-A and HLA-B alleles in patients with COVID-19. **(C-E)** Network analysis of TCR sequences in patients with severe pneumonia and respiratory failure from all diagnosis categories (C), non-COVID-19 groups (D), or COVID-19 (E). Nodes represent unique TCR (CDR3β) sequences and are color-coded by diagnosis. Shared TCR sequences by at least two different pneumonia categories are colored in purple. Edges constitute patterns or specificity groups identified through the GLIPH2 algorithm. **(F-G)** Proportion of HLA-A (F) and HLA-B (G) molecules identified in patients with COVID-19. *n* of patients = 14 **(H-I)** TCRβ (V) gene usage analysis of identified sequences following processing using the GLIPH2 pipeline and cross-matching with the MIRA class I dataset. Proportion of genes by pneumonia category (*n* of patients = non-pneumonia control [4], other pneumonia [7], COVID-19 [14], and other viral pneumonia [8]) (H) and absolute number of TCRβ sequences for each V region by pneumonia category (I) are shown. Dominant genes are annotated in (I).

**Supplemental Figure 14.**
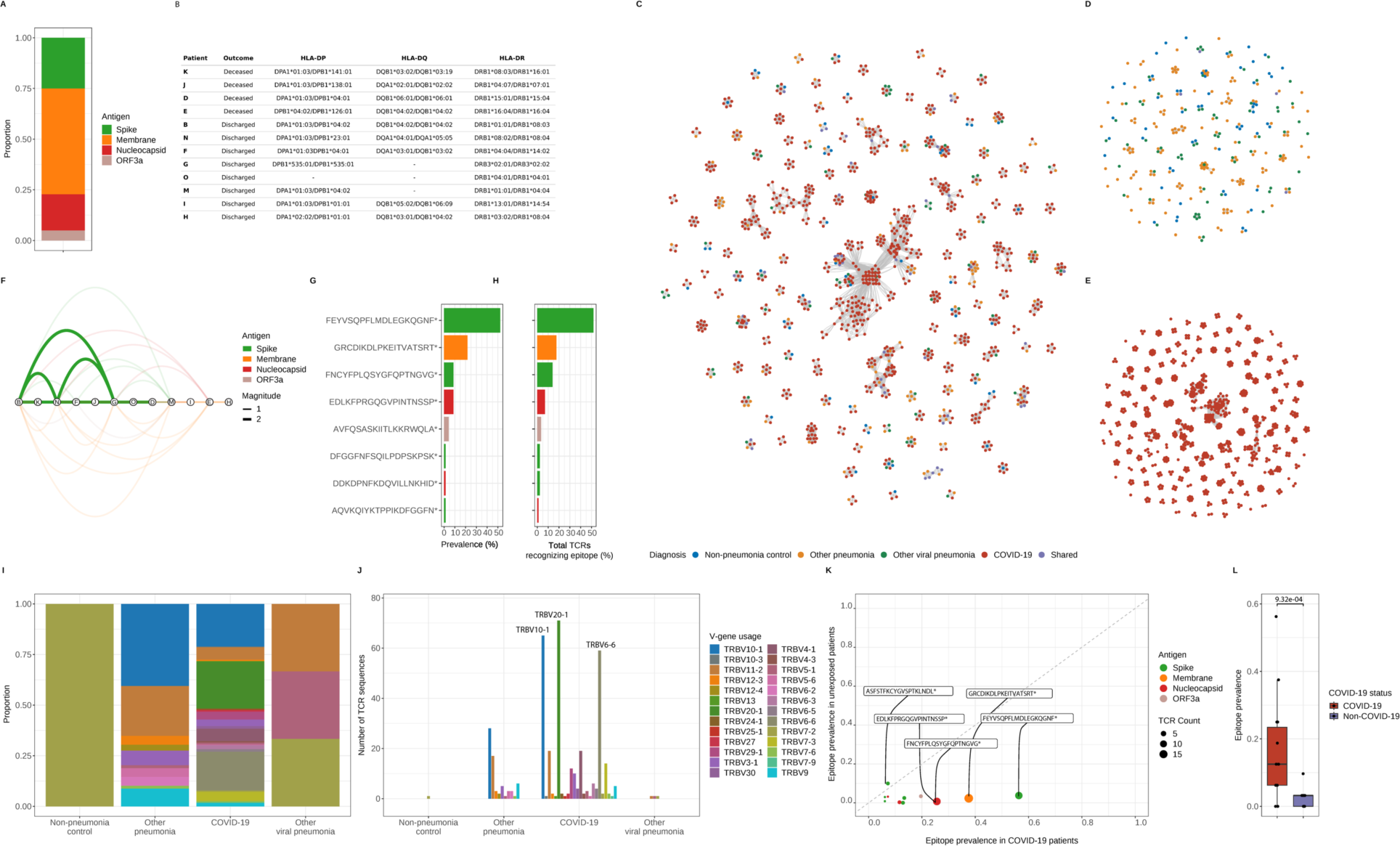
Alveolar CD4^+^ T cell responses during severe SARS-CoV-2 pneumonia. **(A)** Proportion of alveolar CD4^+^ T cell responses by SARS-CoV-2 protein. TCR sequences identified in all samples from patients with COVID-19 were cross-referenced with the MIRA II dataset to identify reactivity against specific SARS-CoV-2 antigens. *n* of patients = 12 and *n* of samples = 22 **(B)** Inferred HLA-DP, HLA-DQ, and HLA-DR alleles in patients with COVID-19. **(C-E)** Network analysis of TCR sequences in patients with severe pneumonia and respiratory failure from all diagnosis categories (C), non-COVID-19 groups (D), or COVID-19 (E). Nodes represent unique TCR (CDR3β) sequences and are color-coded by diagnosis. Shared TCR sequences by at least two different pneumonia categories are colored in purple. Edges constitute patterns or specificity groups identified through the GLIPH2 algorithm. **(F)** Network analysis of shared TCR sequences recognizing SARS-CoV-2 epitopes. Nodes represent unique patients in the COVID-19 group (labeled here using the lettering scheme from Figure 4G), edges constitute shared TCR sequences by at least two patients mapped to a MIRA class II dataset epitope pool, and width of edges (magnitude) denote total number of shared TCR sequences. Edges are color-coded by SARS-CoV-2 antigens. **(G)** Immunoprevalence of SARS-CoV-2 epitopes in patients with COVID-19 was calculated by counting the number of events when a given epitope was shared by at least two patients. Total counts from all eight identified epitopes are represented as percentage (%). **(H)** Overall number of TCR sequences mapped to a given SARS-CoV-2 epitope in patients with COVID-19 was calculated by counting all events of TCRs recognizing an epitope. Total counts from all eight identified epitopes are represented as percentage (%). * denotes other epitopes are present within MIRA class I dataset peptide pool. **(I-J)** TCRβ (V) gene usage analysis of identified sequences following GLIPH2 pipeline and cross-matching with MIRA class II dataset. Proportion of genes by pneumonia category. *n* of patients = non-pneumonia control (1), other pneumonia (4), COVID-19 (12), and other viral pneumonia (2) (I) and absolute number of TCRβ sequences for each V region by pneumonia category (J) are shown. Dominant genes are annotated in (J). **(K)** Scatter plot of SARS-CoV-2 epitope prevalence in patients with COVID-19 (*n* = 12) and without COVID-19 (unexposed, *n* = non-pneumonia control [1], other pneumonia [4], and other viral pneumonia [2]). Dots are color coded by SARS-CoV-2 antigen. Dot size corresponds to the number of detected TCR sequences recognizing a given antigen. **(L)** SARS-CoV-2 epitope prevalence in samples that underwent CD4^+^ TCR sequencing grouped by COVID-19 status. Wilcoxon rank sum test *p*-values are shown.

## Supplemental Data Files

**Supplemental File 1.** Differentially expressed genes in CD8^+^ T cell bulk RNA-sequencing samples from K-means clustering analysis (cluster 1).

**Supplemental File 2.** Differentially expressed genes in CD8^+^ T cell bulk RNA-sequencing samples from K-means clustering analysis (cluster 2).

**Supplemental File 3.** Differentially expressed genes in CD4^+^ T cell bulk RNA-sequencing samples from K-means clustering analysis (cluster 1).

**Supplemental File 4.** Differentially expressed genes in CD4^+^ T cell bulk RNA-sequencing samples from K-means clustering analysis (cluster 2).

**Supplemental File 5.** Significant gene ontology processes identified in CD8^+^ T cell bulk RNA-sequencing samples from K-means clustering analysis (cluster 1).

**Supplemental File 6.** Significant gene ontology processes identified in CD8^+^ T cell bulk RNA-sequencing samples from K-means clustering analysis (cluster 2).

**Supplemental File 7.** Significant gene ontology parent terms and terms identified in CD8^+^ T cell bulk RNA-sequencing samples from K-means clustering analysis (cluster 1).

**Supplemental File 8.** Significant gene ontology parent terms and terms identified in CD8^+^ T cell bulk RNA-sequencing samples from K-means clustering analysis (cluster 2).

**Supplemental File 9.** Significant gene ontology processes identified in CD4^+^ T cell bulk RNA-sequencing samples from K-means clustering analysis (cluster 1).

**Supplemental File 10.** Significant gene ontology parent terms and terms identified in CD4^+^ T cell bulk RNA-sequencing samples from K-means clustering analysis (cluster 1).

**Supplemental File 11.** GLIPH2 analysis establishing specificity groups from CD4^+^ T cell bulk TCR-sequencing samples.

**Supplemental File 12.** GLIPH2 analysis establishing specificity groups from CD8^+^ T cell bulk TCR-sequencing samples.

**Supplemental File 13.** Cross-reference of CD4^+^ T cell repertoire sequences with Multiplex Identification of Antigen-Specific T-Cell Receptors Assay (MIRA) dataset.

**Supplemental File 14.** Cross-reference of CD8^+^ T cell repertoire sequences with Multiplex Identification of Antigen-Specific T-Cell Receptors Assay (MIRA) dataset.

**Supplemental File 15. (A)** Multiplex Identification of Antigen-Specific T-Cell Receptors Assay (MIRA) peptide deconvolution of Class I targets. **(B)** Hierarchical distribution of Class I immunodominant epitopes by patient from the bulk TCR-sequencing subset. **(C)** Pairwise similarity and average conservation score estimation between SARS-CoV-2 and HCoV epitopes.

**Supplemental File 16.** Anonymized metadata by patient and samples for bulk RNA-sequencing and TCR-sequencing analysis.

**Supplemental File 17.** Raw counts for CD4^+^ T cell bulk RNA-sequencing samples.

**Supplemental File 18.** Raw counts for CD8^+^ T cell bulk RNA-sequencing samples.

**Supplemental File 19.** Raw counts for Treg cell bulk RNA-sequencing samples.

**Supplemental File 20.** MiXCR-processed raw sequencing files with TCR repertoire data for CD4^+^ T cell bulk TCR-sequencing samples.

**Supplemental File 21.** MiXCR-processed raw sequencing files with TCR repertoire data for CD8^+^ T cell bulk TCR-sequencing samples.

**Supplemental File 22.** Differential expression analysis in CD8^+^ T cells of pairwise comparison between COVID-19 samples and combined non-COVID-19 samples from Figure 2B.

**Supplemental File 23.** Differential expression analysis in CD4^+^ T cells of pairwise comparison between COVID-19 samples and combined non-COVID-19 samples from Figure 3B.

**Supplemental File 24.** Differential expression analysis in CD8^+^ T cells of pairwise comparison between COVID-19 samples and other viral pneumonia samples from Supplemental Figure 6B

**Supplemental File 25.** Differential expression analysis in CD4^+^ T cells of pairwise comparison between COVID-19 samples and other viral pneumonia samples from Supplemental Figure 7B

**Supplemental File 26.** Differential expression analysis in Treg cells of pairwise comparison between COVID-19 samples and combined non-COVID-19 samples from Supplemental Figure 10B.

**Supplemental File 27.** Differential expression analysis in Treg cells of pairwise comparison between COVID-19 samples and other viral pneumonia samples from Supplemental Figure 10C

**Supplemental File 28.** Differential expression analysis in CD4^+^ T cells of pairwise comparison between early (≤48 hours after intubation) and late (>48 hours after intubation) COVID-19 samples from Supplemental Figure 8A.

**Supplemental File 29.** Differential expression analysis in CD8^+^ T cells of pairwise comparison between early (≤48 hours after intubation) and late (>48 hours after intubation) COVID-19 samples from Supplemental Figure 8B.

**Supplemental File 30.** The NU SCRIPT Study Investigators.

## References

1. Budinger GRS, Misharin AV, Ridge KM, Singer BD, Wunderink RG. Distinctive features of severe SARS-CoV-2 pneumonia. J Clin Invest. 2021;131(14).

2. Grasselli G, et al. Baseline characteristics and outcomes of 1591 patients infected with SARS-CoV-2 admitted to ICUs of the lombardy region, Italy. JAMA. 2020;323(16):1574-1581.

3. Huang C, et al. Clinical features of patients infected with 2019 novel coronavirus in Wuhan, China. Lancet. 2020;395(10223):497–506.

4. Grant RA, et al. Circuits between infected macrophages and T cells in SARS-CoV-2 pneumonia. Nature. 2021;590(7847):635-641.

5. Wendisch D, et al. SARS-CoV-2 infection triggers profibrotic macrophage responses and lung fibrosis. Cell. 2021;184(26):6243–6261 e6227.

6. Flament H, et al. Outcome of SARS-CoV-2 infection is linked to mait cell activation and cytotoxicity. Nat Immunol. 2021;22(3):322–335.

7. Roussel M, et al. Comparative immune profiling of acute respiratory distress syndrome patients with or without SARS-CoV-2 infection. Cell Rep Med. 2021;2(6):100291.

8. Zhang F, et al. IFN-gamma and TNF-alpha drive a cxcl10+ ccl2+ macrophage phenotype expanded in severe COVID-19 lungs and inflammatory diseases with tissue inflammation. Genome Med. 2021;13(1):64.

9. Mathew D, et al. Deep immune profiling of COVID-19 patients reveals distinct immunotypes with therapeutic implications. Science. 2020;369(6508).

10. Kuri-Cervantes L, et al. Comprehensive mapping of immune perturbations associated with severe COVID-19. Sci Immunol. 2020;5(49).

11. Moss P. The T cell immune response against SARS-CoV-2. Nat Immunol. 2022;23(2):186–193.

12. Rydyznski Moderbacher C, et al. Antigen-specific adaptive immunity to SARS-CoV-2 in acute COVID-19 and associations with age and disease severity. Cell. 2020;183(4):996–1012 e1019.

13. Lucas C, et al. Longitudinal analyses reveal immunological misfiring in severe COVID-19. Nature. 2020;584(7821):463-469.

14. Pickens CO, et al. Bacterial superinfection pneumonia in patients mechanically ventilated for COVID-19 pneumonia. Am J Respir Crit Care Med. 2021;204(8):921–932.

15. Gao CA, et al. Machine learning links unresolving secondary pneumonia to mortality in patients with severe pneumonia, including COVID-19. J Clin Invest. 2023;133(12).

16. Markov NS, et al. SCRIPT CarpeDiem dataset: Demographics, outcomes, and per-day clinical parameters for critically ill patients with suspected pneumonia (version 1.1.0). 2023. 10.13026/5phr-4r89.

17. Recovery Collaborative Group, et al. Dexamethasone in hospitalized patients with Covid-19. N Engl J Med. 2021;384(8):693–704.

18. Singh G, et al. Low BALF CD4 T cells count is associated with extubation failure and mortality in critically ill covid-19 pneumonia. Ann Med. 2022;54(1):1894–1905.

19. Szabo PA, et al. Longitudinal profiling of respiratory and systemic immune responses reveals myeloid cell-driven lung inflammation in severe COVID-19. Immunity. 2021;54(4):797–814 e796.

20. Martin TR. Neutrophils and lung injury: Getting it right. J Clin Invest. 2002;110(11):1603–1605.

21. Pickens CI, et al. An adjudication protocol for severe pneumonia. Open Forum Infect Dis. 2023;10(7):ofad336.

22. Bergamaschi L, et al. Longitudinal analysis reveals that delayed bystander CD8+ T cell activation and early immune pathology distinguish severe COVID-19 from mild disease. Immunity. 2021;54(6):1257–1275 e1258.

23. Neidleman J, et al. Distinctive features of SARS-CoV-2-specific T cells predict recovery from severe COVID-19. Cell Rep. 2021;36(3):109414.

24. Le Bert N, et al. Highly functional virus-specific cellular immune response in asymptomatic SARS-CoV-2 infection. J Exp Med. 2021;218(5).

25. Tan AT, et al. Early induction of functional SARS-CoV-2-specific T cells associates with rapid viral clearance and mild disease in COVID-19 patients. Cell Rep. 2021;34(6):108728.

26. Westblade LF, et al. SARS-CoV-2 viral load predicts mortality in patients with and without cancer who are hospitalized with COVID-19. Cancer Cell. 2020;38(5):661–671 e662.

27. Pujadas E, et al. SARS-CoV-2 viral load predicts COVID-19 mortality. Lancet Respir Med. 2020;8(9):e70.

28. Wolfel R, et al. Virological assessment of hospitalized patients with COVID-2019. Nature. 2020;581(7809):465-469.

29. Jovisic M, Mambetsariev N, Singer BD, Morales-Nebreda L. Differential roles of regulatory T cells in acute respiratory infections. J Clin Invest. 2023;133(14).

30. Zhao J, et al. Airway memory CD4(+) T cells mediate protective immunity against emerging respiratory coronaviruses. Immunity. 2016;44(6):1379–1391.

31. Woodland DL, Blackman MA. Immunity and age: Living in the past? Trends Immunol. 2006;27(7):303–307.

32. Goronzy JJ, Fang F, Cavanagh MM, Qi Q, Weyand CM. Naive T cell maintenance and function in human aging. J Immunol. 2015;194(9):4073–4080.

33. Goronzy JJ, Weyand CM. Successful and maladaptive T cell aging. Immunity. 2017;46(3):364–378.

34. Huang H, Wang C, Rubelt F, Scriba TJ, Davis MM. Analyzing the mycobacterium tuberculosis immune response by T-cell receptor clustering with gliph2 and genome-wide antigen screening. Nat Biotechnol. 2020;38(10):1194–1202.

35. Lineburg KE, et al. CD8(+) T cells specific for an immunodominant SARS-CoV-2 nucleocapsid epitope cross-react with selective seasonal coronaviruses. Immunity. 2021;54(5):1055–1065 e1055.

36. Pogorelyy MV, et al. Resolving SARS-CoV-2 CD4(+) T cell specificity via reverse epitope discovery. Cell Rep Med. 2022;3(8):100697.

37. Nolan S, et al. A large-scale database of T-cell receptor beta (tcrbeta) sequences and binding associations from natural and synthetic exposure to SARS-CoV-2. Res Sq. 2020;10.21203/rs.3.rs-51964/v1.

38. Moutaftsi M, et al. Uncovering the interplay between CD8, CD4 and antibody responses to complex pathogens. Future Microbiol. 2010;5(2):221-239.

39. Torres Acosta MA, Singer BD. Pathogenesis of COVID-19-induced ARDS: Implications for an ageing population. Eur Respir J. 2020;56(3).

40. Onder G, Rezza G, Brusaferro S. Case-fatality rate and characteristics of patients dying in relation to COVID-19 in Italy. JAMA. 2020;323(18):1775–1776.

41. Peters B, Nielsen M, Sette A. T cell epitope predictions. Annu Rev Immunol. 2020;38:123–145.

42. Orenbuch R, Filip I, Comito D, Shaman J, Pe’er I, Rabadan R. arcasHLA: High-resolution HLA typing from RNAseq. Bioinformatics. 2020;36(1):33–40.

43. Reynisson B, Alvarez B, Paul S, Peters B, Nielsen M. NetMHCpan-4.1 and netmhciipan-4.0: Improved predictions of MHC antigen presentation by concurrent motif deconvolution and integration of ms MHC eluted ligand data. Nucleic Acids Res. 2020;48(W1):W449–W454.

44. Gonzalez-Galarza FF, et al. Allele frequency net database (AFND) 2020 update: Gold-standard data classification, open access genotype data and new query tools. Nucleic Acids Res. 2020;48(D1):D783–D788.

45. Sidney J, Peters B, Sette A. Epitope prediction and identification-adaptive T cell responses in humans. Semin Immunol. 2020;50:101418.

46. Le Bert N, et al. SARS-CoV-2-specific T cell immunity in cases of COVID-19 and SARS, and uninfected controls. Nature. 2020;584(7821):457-462.

47. Grifoni A, et al. Targets of T cell responses to SARS-CoV-2 coronavirus in humans with COVID-19 disease and unexposed individuals. Cell. 2020;181(7):1489–1501 e1415.

48. Sekine T, et al. Robust T cell immunity in convalescent individuals with asymptomatic or mild COVID-19. Cell. 2020;183(1):158–168 e114.

49. Bacher P, et al. Low-avidity CD4(+) T cell responses to SARS-CoV-2 in unexposed individuals and humans with severe COVID-19. Immunity. 2020;53(6):1258–1271 e1255.

50. Weiskopf D, et al. Phenotype and kinetics of SARS-CoV-2-specific T cells in COVID-19 patients with acute respiratory distress syndrome. Sci Immunol. 2020;5(48).

51. Dykema AG, et al. Functional characterization of CD4+ T cell receptors crossreactive for SARS-CoV-2 and endemic coronaviruses. J Clin Invest. 2021;131(10).

52. Mateus J, et al. Selective and cross-reactive SARS-CoV-2 T cell epitopes in unexposed humans. Science. 2020;370(6512):89-94.

53. Schulien I, et al. Characterization of pre-existing and induced SARS-CoV-2-specific CD8(+) T cells. Nat Med. 2021;27(1):78–85.

54. Sun X, et al. Longitudinal analysis reveals age-related changes in the T cell receptor repertoire of human T cell subsets. J Clin Invest. 2022;132(17).

55. Morales-Nebreda L, et al. Aging imparts cell-autonomous dysfunction to regulatory T cells during recovery from influenza pneumonia. JCI Insight. 2021;6(6).

56. Gao CA, Morales-Nebreda L, Pickens CI. Gearing up for battle: Harnessing adaptive T cell immunity against gram-negative pneumonia. Front Cell Infect Microbiol. 2022;12:934671.

57. Singer BD, Jain M, Budinger GRS, Wunderink RG. A call for rational Intensive care in the era of COVID-19. Am J Respir Cell Mol Biol. 2020;10.1165/rcmb.2020-0151LE.

58. Channappanavar R, et al. Dysregulated type I interferon and inflammatory monocyte-macrophage responses cause lethal pneumonia in SARS-CoV-infected mice. Cell Host Microbe. 2016;19(2):181–193.

59. Zhao J, Zhao J, Perlman S. T cell responses are required for protection from clinical disease and for virus clearance in severe acute respiratory syndrome coronavirus-infected mice. J Virol. 2010;84(18):9318–9325.

60. McMahan K, et al. Correlates of protection against SARS-CoV-2 in rhesus macaques. Nature. 2021;590(7847):630-634.

61. Cameron MJ, et al. Interferon-mediated immunopathological events are associated with atypical innate and adaptive immune responses in patients with severe acute respiratory syndrome. J Virol. 2007;81(16):8692–8706.

62. Hadjadj J, et al. Impaired type I interferon activity and inflammatory responses in severe COVID-19 patients. Science. 2020;369(6504):718-724.

63. Rha MS, et al. PD-1-expressing SARS-CoV-2-specific CD8(+) T cells are not exhausted, but functional in patients with COVID-19. Immunity. 2021;54(1):44–52 e43.

64. Bertoletti A, Tan AT, Le Bert N. The T-cell response to SARS-CoV-2: Kinetic and quantitative aspects and the case for their protective role. Oxford Open Immunology. 2021;2(1).

65. Saini SK, et al. SARS-CoV-2 genome-wide T cell epitope mapping reveals immunodominance and substantial CD8(+) T cell activation in COVID-19 patients. Sci Immunol. 2021;6(58).

66. Ng KW, et al. Preexisting and de novo humoral immunity to SARS-CoV-2 in humans. Science. 2020;370(6522):1339-1343.

67. Braun J, et al. SARS-CoV-2-reactive T cells in healthy donors and patients with COVID-19. Nature. 2020;587(7833):270-274.

68. Bartolo L, et al. SARS-CoV-2-specific T cells in unexposed adults display broad trafficking potential and cross-react with commensal antigens. Sci Immunol. 2022;7(76):eabn3127.

69. Pothast CR, et al. SARS-CoV-2-specific CD4(+) and CD8(+) T cell responses can originate from cross-reactive cmv-specific T cells. eLife. 2022;11.

70. van Diepen S, McAlister FA, Chu LM, Youngson E, Kaul P, Kadri SS. Association between vaccination status and outcomes in patients admitted to the ICU with COVID-19. Crit Care Med. 2023;51(9):1201–1209.

71. Starren JB, Winter AQ, Lloyd-Jones DM. Enabling a learning Health system through a unified enterprise data warehouse: The experience of the northwestern university clinical and translational sciences (NUCATS) institute. Clin Transl Sci. 2015;8(4):269–271.

72. Walter JM, Helmin KA, Abdala-Valencia H, Wunderink RG, Singer BD. Multidimensional assessment of alveolar T cells in critically ill patients. JCI Insight. 2018;3(17):e123287.

73. Acute Respiratory Distress Syndrome Network, et al. Ventilation with lower tidal volumes as compared with traditional tidal volumes for acute lung injury and the acute respiratory distress syndrome. N Engl J Med. 2000;342(18):1301-1308.

74. Guerin C, et al. Prone positioning in severe acute respiratory distress syndrome. N Engl J Med. 2013;368(23):2159–2168.

75. Combes A, et al. Extracorporeal membrane oxygenation for severe acute respiratory distress syndrome. N Engl J Med. 2018;378(21):1965–1975.

76. Kurihara C, et al. Extracorporeal membrane oxygenation can successfully support patients with severe acute respiratory distress syndrome in lieu of mechanical ventilation. Crit Care Med. 2018;46(11):e1070–e1073.

77. Singer BD, Corbridge TC. Pressure modes of invasive mechanical ventilation. South Med J. 2011;104(10):701–709.

78. Beigel JH, et al. Remdesivir for the treatment of Covid-19 - final report. N Engl J Med. 2020;383(19):1813–1826.

79. Ewels PA, et al. The nf-core framework for community-curated bioinformatics pipelines. Nat Biotechnol. 2020;38(3):276–278.

80. Di Tommaso P, Chatzou M, Floden EW, Barja PP, Palumbo E, Notredame C. Nextflow enables reproducible computational workflows. Nat Biotechnol. 2017;35(4):316–319.

81. Dobin A, et al. STAR: Ultrafast universal RNA-seq aligner. Bioinformatics. 2013;29(1):15–21.

82. Patro R, Duggal G, Love MI, Irizarry RA, Kingsford C. Salmon provides fast and bias-aware quantification of transcript expression. Nat Methods. 2017;14(4):417–419.

83. Robinson MD, McCarthy DJ, Smyth GK. edgeR: A Bioconductor package for differential expression analysis of digital gene expression data. Bioinformatics. 2010;26(1):139–140.

84. Gu Z, Eils R, Schlesner M. Complex heatmaps reveal patterns and correlations in multidimensional genomic data. Bioinformatics. 2016;32(18):2847–2849.

85. Rahnenfuhrer AA. topGO: Enrichment analysis for gene ontology. 2023. 10.18129/B9.bioc.topGO.

86. Durinck S, Spellman PT, Birney E, Huber W. Mapping identifiers for the integration of genomic datasets with the R/Bioconductor package biomart. Nat Protoc. 2009;4(8):1184–1191.

87. Sayols S. Rrvgo: A Bioconductor package for interpreting lists of gene ontology terms. MicroPubl Biol. 2023;2023.

88. Subramanian A, et al. Gene set enrichment analysis: A knowledge-based approach for interpreting genome-wide expression profiles. Proc Natl Acad Sci U S A. 2005;102(43):15545–15550.

89. Korotkevich G, Sukhov V, Sergushichev A. Fast gene set enrichment analysis. bioRxiv. 2019;10.1101/060012.060012.

90. Aliee H, Theis FJ. AutoGeneS: Automatic gene selection using multi-objective optimization for RNA-seq deconvolution. Cell Syst. 2021;12(7):706–715 e704.

91. Bolotin DA, et al. Mixcr: Software for comprehensive adaptive immunity profiling. Nat Methods. 2015;12(5):380–381.

92. Nazarov V, et al. immunarch: Bioinformatics analysis of T-cell and B-cell immune repertoires. 2023. 10.5281/zenodo.3367200.

93. Butts C. network: Classes for relational data. 2015. https://CRAN.R-project.org/package=network

94. Pedersen T. tidygraph: A tidy API for graph manipulation. 2023. https://tidygraph.data-imaginist.com

95. Pedersen T. ggraph: An implementation of grammar of graphics for graphs and networks. 2022, https://ggraph.data-imaginist.com, https://github.com/thomasp85/ggraph.

96. Ihaka R, Gentleman R. R: A language for data analysis and graphics. Journal of Computational and Graphical Statistics. 1996;5:299–314.

97. Constantin A, Patil I. ggsignif: R package for displaying significance brackets for ‘ggplot2’. PsyArxiv. 2021;doi:10.31234/osf.io/7awm6.

98. Kassambara A. ggpubr: ‘ggplot2’ based publication ready plots. 2023. https://cran.r-project.org/web/packages/ggpubr/index.html

99. Wickham H ggplot2: Elegant graphics for data analysis. New York: Springer-Verlag; 2016.

